# Nkx2.5-dependent alterations of the embryonic heart DNA methylome identify novel cis-regulatory elements in cardiac development

**DOI:** 10.1101/186395

**Authors:** Bushra Gorsi, Timothy L. Mosbruger, Megan Smith, Jonathon T. Hill, H. Joseph Yost

## Abstract

Transcription factor Nkx2.5 is frequently mutated in congenital heart disease, but the mechanisms by which Nkx2.5 regulates heart development are poorly understood. By generating comprehensive DNA methylome maps from zebrafish embryonic hearts in *nxk2.5* mutants and siblings, we discovered that Nkx2.5 regulates DNA methylation patterns during cardiac morphogenesis. We identified hundreds of Nkx-dependent heart-specific Differentially Methylated Regions (nhDMRs). A majority of the nhDMRs were hypomethylated in *nkx2.5*^−/-^ hearts, correlating with changes in the mutant transcriptome, suggesting Nkx2.5 functions largely as a repressor. Distinct Nkx DNA-binding motifs were significantly enriched in subclasses of nhDMRs. Furthermore, nhDMRs were significantly associated with histone H3K4me1 and H3K27ac post-translational modifications, suggesting Nkx2.5 regulates gene expression by differential methylation of cis-regulatory elements. Using transgenics, we validated several nhDMRs with enhancer activities in the heart. We propose a novel role of Nkx2.5 mediated DNA methylation is integral in activating and repressing Nkx2.5 target genes during heart development.

## Introduction

Nkx2.5 is one of the most commonly mutated transcription factors in human Congenital Heart Disease (CHD), occurring in 1-4% of cardiac malformations (1). Nkx2.5 is a dual activator and repressor required for cardiac progenital cell (CPC) specification, chamber growth, and formation and maintenance of the conduction system (2). Although its function is critical during early cardiomyogenesis, our understanding of how Nkx2.5 can both activate and repress gene function is limited. In zebrafish, Nkx2.5 has a role in heart tube extension and chamber morphogenesis by maintaining normal ratios of ventricular and atrial cells (3,4). The ratio of atrial to ventricle cells is skewed to significantly more atrial cells, and the ventricle is diminished. Although *nkx2.5* is not necessary to induce ventricle initiation, it is required for the maintenance of ventricle identity from early to late stages of heart development (5). In the absence of *nkx2.5*, ventricular cells transdifferentiate to atrial cells (3), suggesting Nkx2.5 is functioning to repress the atrial gene program to maintain ventricle identity. Nkx2.5 also has been shown to be an important mediator of endothelial cell differentiation in pharyngeal arch (PAA) development, as Nkx2.5 actively represses angioblast differentiation early in the anterior lateral plate mesoderm but is required for PAA angioblast differentiation in later stages (6). The dual role of Nkx2.5 to function as a transcriptional activator and repressor in early lineage commitment decisions is not unique to zebrafish, but is also conserved in mice. Partial reduction or a complete loss of Nkx2.5 from an *Nkx2.5* hypomorphic mouse heart and Nkx2.5 knockout in ESC-derived cardiomyocytes, showed a large proportion of differentially genes are upregulated and have Nkx2.5 binding motifs at their loci, lending support to the hypothesis that Nkx2.5 functions as a repressor (7,8). In mouse ESC-derived Nkx2.5 knockout cardiomyocytes, CPCs differentiate quicker than the wildtype CPC, suggesting Nkx2.5 may act early to repress differentiation genes, such as Bmp2/Smad1, to maintain the CPC population (8,9). During stages of cardiac maturation in *Nkx2.5* hymorphic mice, there is a direct repression of cardiac fast troponins isoforms in the atrium, but not the ventricle, another example of Nkx2.5 mediated lineage specific repression (7). It is well established that Nkx2.5 cooperativity with other transcription factors such as Tbx5, Gata4, Tbx2, Tbx20, SRF and Meis1 facilitates its activating and repressing functions (7,10–12). This largely occurs through the conserved homeodomain, which serves as both a sequence-specific DNA binding domain and a protein-protein interaction domain. Molecular analysis of the homeodomain suggest a key regulatory role of Nkx2.5 in repressing gene functions through the direct interaction and/or recruitment of epigenetic modifiers such as Histone deacetylases (HDACs) that deacetylate histone (12). However HDACs are thought to not function alone and operate with pathways that involve DNA methyltransferases (DNMTs). Nkx2.5 can also repress transcription of target genes through direct interaction with histone H3K36me3 methyltransferase (13), this histone mark has also been shown to serve as a mark for HDACs (14) and DNMT3B (15). Moreover, DNMTs can recruit HDACs directly, suggesting methylation and deacetylation act together to establish a repressed state (16,17).

Direct epigenetic configuration of chromatin might be one of the mechanisms by which Nkx2.5 can function as a dual activator and repressor in embryonic heart development. We sought to address whether Nkx2.5 might serve as a functional repressor through the evolutionary conserved process of DNA methylation. Methylation of cytosines is thought to prevent gene transcription by directly interfering with the recognition sequence of some TFs, while specifically binding other TFs. Emerging studies have identified homeobox proteins including the Nkx family of proteins (Nkx2.2 and Nkx2.5) as interacting with methylated DNA via their homeodomian as well as binding to methylated CpGs in TF binding motif (18–20). Thus analysing the DNA methylome in Nkx2.5 mutants will provide a gateway for studying the interactions of Nkx2.5 with the DNA methylome.

Using a genomics approach, we generated comprehensive DNA methylome maps and RNA-seq profiles from isolated 48hpf zebrafish hearts and non-heart tissue from both *nkx2.5*^−/-^ mutants and siblings. We discovered thousands of <u>N</u>kx2.5-dependent <u>h</u>eart-specific <u>D</u>ifferentially <u>M</u>ethylated <u>R</u>egions (nhDMRs), and found significant enrichment of the nhDMRs at distal cis-regulatory regions. A majority of the nhDMRs were hypomethylated in the *nkx2.5*^−/-^ hearts, positively correlating with a large number of differentially upregulated genes in *nkx2.5*^−/-^ mutants, corroborating the idea that Nkx2.5 acts largely as a repressor of gene function. nhDMRs neighboring the differentially genes were enriched in Nkx motifs. Our data shows for the first time a class of cardiac-specific distal and proximal regulatory elements in zebrafish that are dependent on Nkx2.5 regulation, and tests a selection of these elements for their ability to regulate cardiac expression in stable transgenic lines. The hundreds of Nkx2.5 dependent regulatory elements will serve as a resource for further functional exploration of coding and non-coding elements that contribute to cardiac disease. Furthermore, we propose that Nkx2.5 imparts a level of regulation by altering DNA methylation patterns in the heart.

## RESULTS

### *nkx2.5* mutant hearts have differentially methylated DNA regions compared to sibling hearts

We performed whole genome-wide Bisulfite-seq and RNA-seq analysis at 48 hours post fertilization (hpf) to identify changes in the DNA methylome and transcriptome of *nkx2.5*^−/-^ mutants that might contribute to the abberant heart development. We manually dissected *nkx2.5*^−/-^ heart tissue and sibling heart tissue from whole embryos and collected DNA from three replicates of the following four samples: *nkx2.5*^−/-^ heart, *nkx2.5*^−/-^ body, sib heart and sib body. DNA samples had a successful bisulfite conversion rate of 99.9%. Using BisSeq (Useq v.8.8.5 package) we analyzed the changes in DNA methylation in *nkx2.5*^−/-^ heart versus sib heart, compared to mutant and wildtype non-heart tissue. We imposed quality filters of ≥8 reads in a window and 5 ≥ CpGs in a region. Using a cutoff criterion of 1.5-fold change and p< 0.05, we found 3702 unique differentially methylated regions (DMRs) with 914 hypermethylated DMRs in the *nkx2.5*^−/-^ heart and 2788 hypomethylated DMRs in the *nkx2.5*^−/-^ heart (Figure 1A). Nkx dependent heart-specific DMRs (nhDMRs) were defined as DMRs that did not overlap with DMRs observed in the *nkx2.5*^−/-^ heart v *nkx2.5*^−/-^ body and sib heart v sib body comparisons. A large proportion of the nhDMRs were located within 10kb of the nearest TSS for both hypermethylated (41%) and hypomethylated (37%) nhDMRs (Figure 1B) accompanied with the highest log2ratio fold change and percent changes in DNA methylation (Figure 1C). A moderate percent change in nhDMRs was observed at short and long range distances to TSS. By comparison, percent changes in DMRs observed in the *nkx2.5*^−/-^ heart v *nkx2.5*^−/-^ body and sib heart v sib body were smaller and a shorter range (Figure S1A-B). Average length of subclasses of *nkx2.5*^−/-^ nhDMRs was 250bp, similar to DMRs in non-heart tissue (Figure 1D).

**Figure 1:**
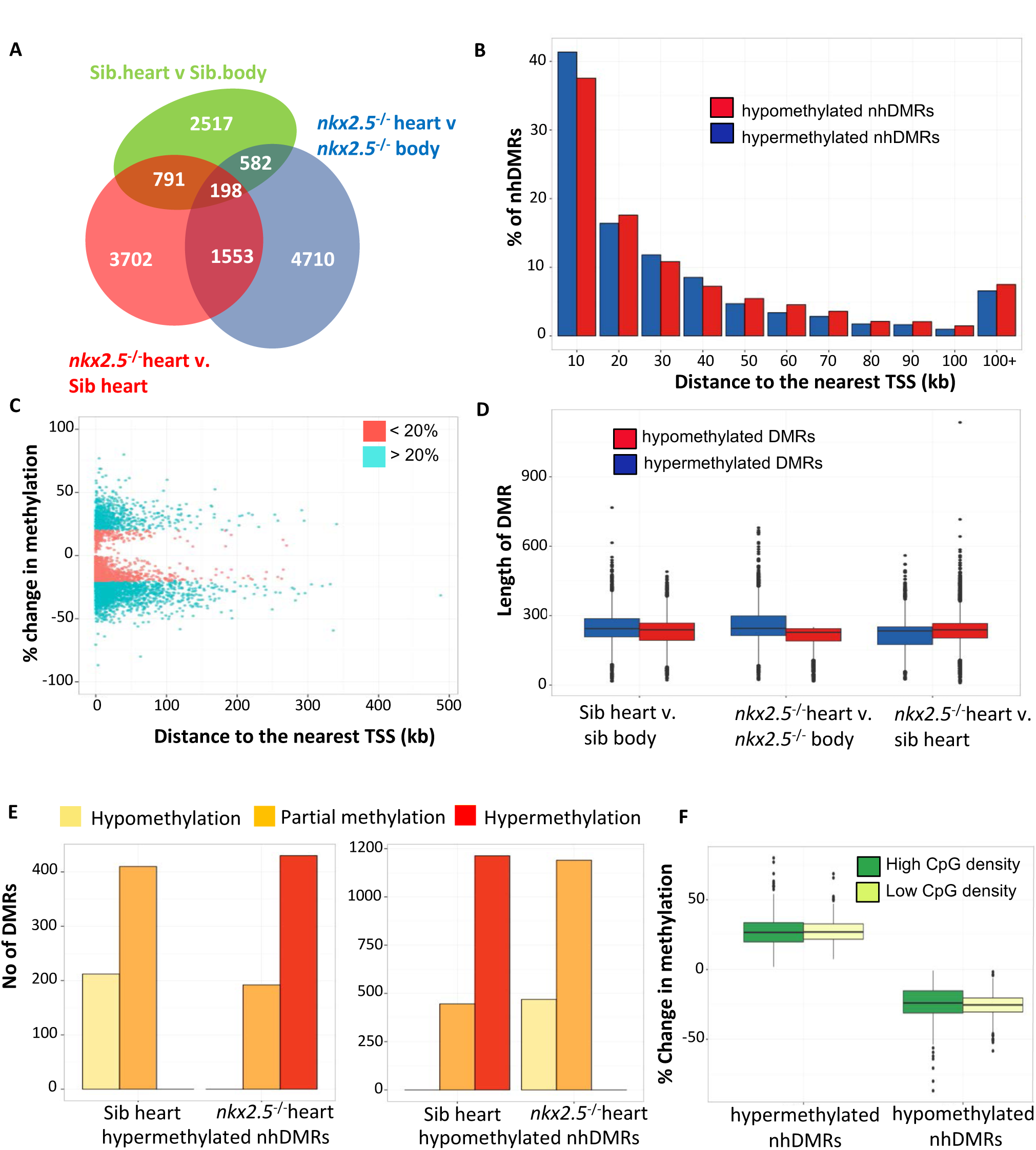
Distribution of nkx2.5-dependent heart-specific differentially methylated regions (nhDMRs). (A) Venn diagram of DMRs in *nkx2.5*^−/-^ heart compared to *nkx2.5*^−/-^ body, sib heart and sib body. A larger number of DMRs are present (4710) in *nkx2.5*^−/-^ heart when comparing to *nkx2.5*^−/-^ non-heart tissue versus sib heart (3702) DMRs. DMRs that overlapped by 1bp or more were consolidated and redundancies removed from further analysis. (B) Percentage distribution of hypomethylated (red) and hypermethylated (blue) nhDMRs over distance to the nearest TSS. (C) Percent changes in methylation of nhDMRs from *nkx2.5*^−/-^ v. sib heart plotted against distance to the nearest TSS. (D) Length of DMRs in mutants and siblings, heart and non-heart tissue have a median range of 250bp, comparable across different genotypes and tissue types. (E). nhDMRs in *nkx2.5*^−/-^ *v* sib heart were categorized into hypermethylation (≥0.7), partial methylation (≥0.3 to <0.7) and hypomethylation (<0.3). The categories were designated based on tissue level expression of the known heart and non heart genes. (F) Hypermethylated and hypomethylated nhDMRs with the greatest changes in methylation are categorized comparably into high and low CpG density nhDMRs.

We next examined the nhDMRs in the context of hypomethylation, partial methylation and hypermethylation in the *nkx2.5*^−/-^ heart. In order to define hypo and hyper methylation boundaries, we examined genes that encode cardiac transcription factors, ion channels, endocardial and epicardial genes that are expressed or silenced in our *nkx2.5*^−/-^ RNA-seq dataset (summarized in heatmaps, Figure S2A-B). Methylation status of these genes showed negative correlation with gene expression. Therefore, we took the average fraction methylation at the TSS and defined hypomethylated regions as ≤0.3, partially methylated regions as ranging from ≥0.3 to ≤0.7, and hypermethylated regions as ≥0.7. Percentage of DNA methylation of individual CpGs showed clear separation between hypomethylation, hypermethylation and partial methylation (Figure 1E) of DMRs between *nkx2.5*^−/-^ heart and sib hearts. High percent changes of methylation were associated with both classes of nhDMRs with high and low CpG densities, suggesting nhDMRs are not exclusively at promoters (Figure 1F).

### Distribution of DMRs show significant enrichment at the promoter in *nkx2.5* mutant heart

We next examined the genomic locations of *nkx2.5*-dependent nhDMRs in *nkx2.5*^−/-^ mutant hearts. We defined filtered nhDMRs as nhDMRs that had greater than 20% methylation change. Using the Cis-regulatory Element Annotation System (CEAS), we found that genomic distribution of filtered nhDMRs was highly nonrandom. The largest percentage of filtered nhDMRs were observed in intergenic and intronic regions. We found a significantly lower percentage of intronic nhDMRs, but a greater percentage of intergenic nhDMRs than expected by random chance (Figure 2A, red asterisk). Approximately 2.6% of hypomethylated nhDMRs and 2.0% of hypermethylated nhDMRs were within the proximal promoter, greater than predicted by chance distribution throughout the genome (Figure 2B). The data suggests that promoters were enriched as main targets of DNA methylation, as one would expect since CpG islands are largely located at the TSS and that regulations of nhDMRs is likely to occur proximal to the promoters of genes.

**Figure 2:**
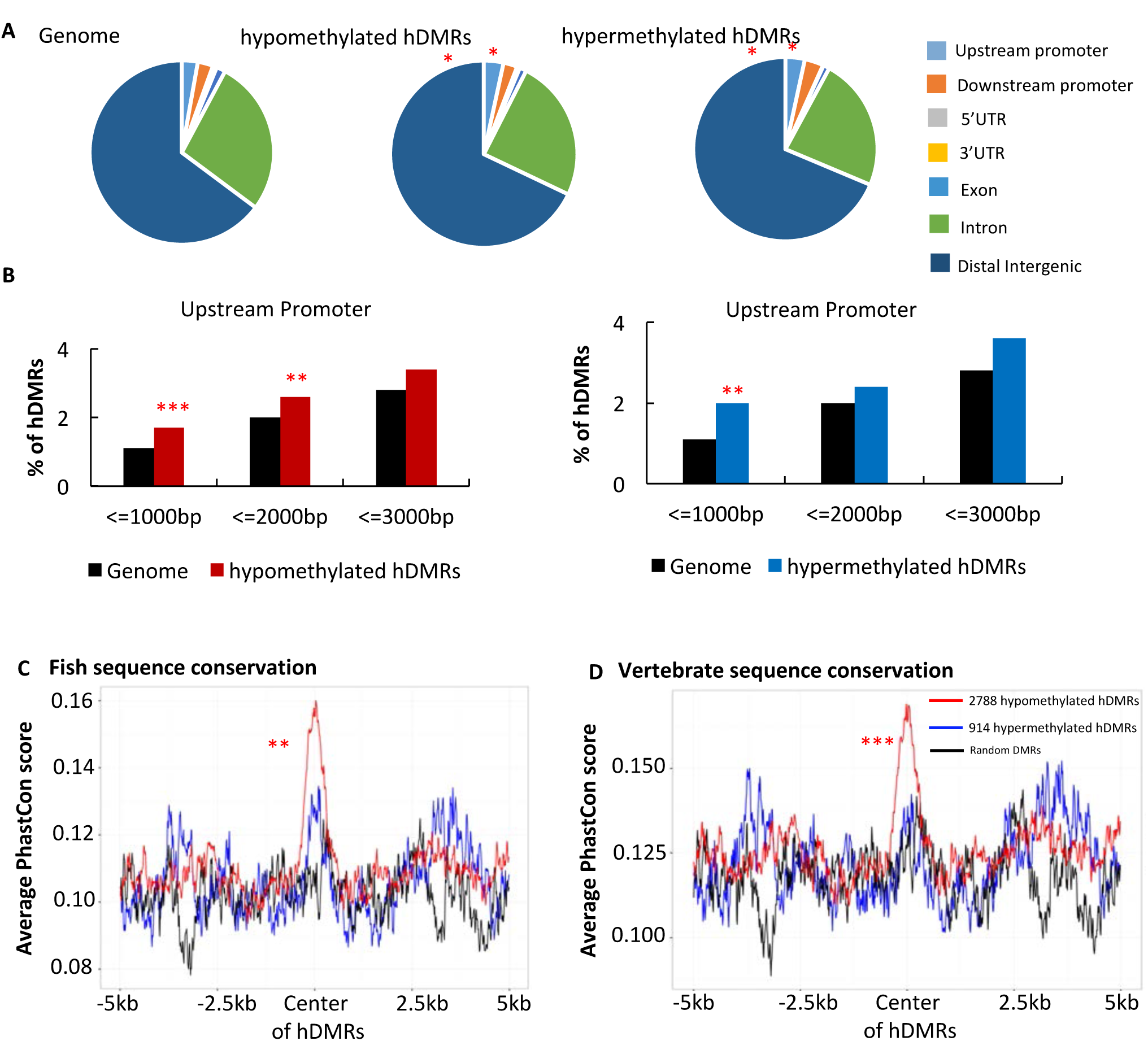
Genomic distribution *nkx2.5*-dependent hypomethylated and hypermethylated nhDMRs. (A) Assignment of nhDMRs to genomic regions indicates increased representation in upstream promoters and distal intergenic regions. (B) Overrepresentation of nhDMRs in upstream promoter regions. Significant enrichment of the nhDMRs at the promoter in both hypermethylated and hypomethylated DMRs in the *nkx2.5*^−/-^ heart. * p < 0.01 ** p < 0.001, *** p <0.0001 (C, D) Sequence conservation of hypomethylated and hypermethylated nhDMRs in (C) multiple fish species and (D) other veterbrates; average phastcon scores were plotted over 10kb of regions centered on hypomethylated nhDMRs (red), hypermethylated nhDMRs (blue), compared to random (black) DMRs.

### Nkx2.5-dependent nhDMRs are highly conserved in vertebrates

To test whether both classes of nhDMRs, hypermethylated and hypomethylated, were evolutionarily conserved we obtained the average PhastCon scores based on an eight-way fish and vertebrate genome alignment with zebrafish using the UCSC browser. We found higher conservation scores for both hypermethylated and hypomethylated nhDMRs than for neighboring regions when aligned to multiple fish genomes (Figure 2C) and multiple vertebrate genomes (Figure 2D). In particular, a significant proportion of hypomethylated nhDMRs contained sequences that were evolutionarily conserved across vertebrates and fish (permutation test, p-value <0.001). The high level of significant conservation of sequences at the center of nhDMRs suggests these nhDMRs have functional significance across vertebrates, and will serve as a valuable resource for the searches in human whole genomes for non-coding variants that might be causative of CHD. However, lack of conservation in an nhDMRs does not rule out its regulatory function (21).

### Subsets of both classes of nhDMRs show regulatory function

To evaluate the functional potential of both classes of nhDMRs (hypomethylated and hypermethylated), we intersected our data with 48hpf whole embryo ChIP-seq data enriched for the following active histone post-translational modifications (PTMs): H3K4me1, H3K4me3 and H3K27ac(22). Distal and proximal regulatory elements have the capability of turning on gene expression independent of location and orientation relative to the gene of interest. In turn, this makes identifying distal and proximal regulatory elements particularly challenging. However, various experimental and computational approaches have identified signature histone PTMs, H3K4me1 and H3K27ac, that define regulatory elements (23–25). Using these criteria, we determined whether the nhDMRs were correlated with regulatory elements of nearby genes. Strikingly, we observed high ChIP-seq signals for H3K4me1, H3K4me3 and H3K27ac PTMs centered over hypomethylated nhDMRs (Figure 3A,C,E). More than half of the hypomethylated nhDMRs intersected with single, double and multiple active histone PTMs with a significant number associated with the triple histone H3K4me3; H3K4me1; H3K27ac (Fisher test, p-value <0.001) and double histone H3K4me1; H3K27ac active PTMs (Fisher test, p-value < 0.001), indicating these nhDMRs elements are correlated with active regulatory function. Conversely, the H3K4me3 signal was lower at the center of hypermethylated nhDMRs than in neighboring regions, but there was increased H3K4me1 and H3K27ac PTMs centered at the hypermethylated nhDMRs (Figure 3B,D). We identified a significant number of hypermethylated nhDMRs also associated with double histone PTMs (H3K4me1; H3K27ac, Fisher test, p-value 5.55 x10^−84^) (Figure 3F). Considering the evolutionary conservation of both classes of nhDMRs and the striking integration with histone PTM patterns, we identified a number of heart regulatory elements that are differentially methylated in *nkx2.5*^−/-^ heart.

**Figure 3:**
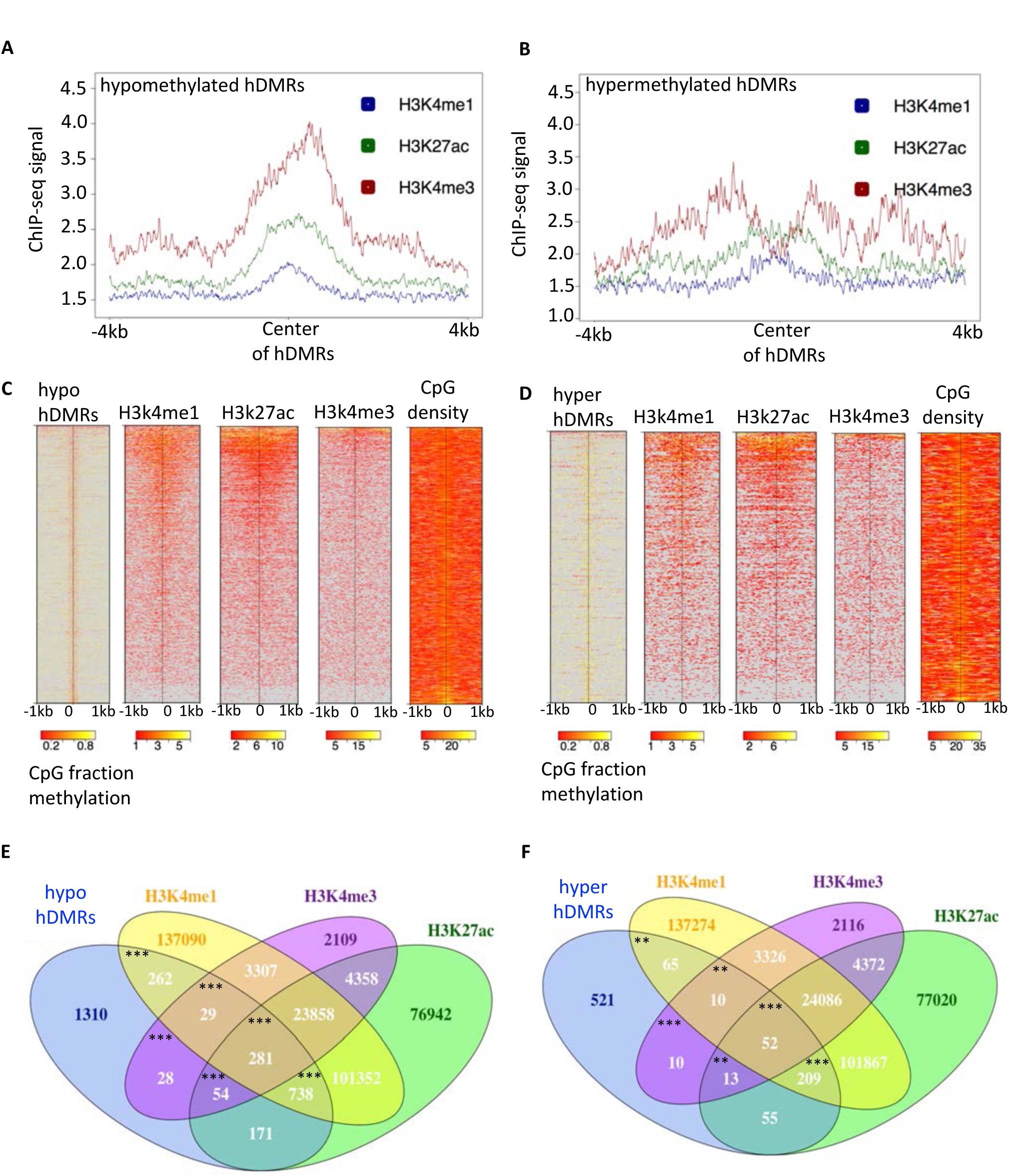
nhDMRs significantly associate with histone PTM marks representative of enhancer elements. Averaged ChIP-seq signals of active histone PTM marks from 48hpf embryos were plotted over +/-4kb regions centered on (A) hypomethylated nhDMRs and (B) hypermethylated nhDMRs. (C, D) Heatmaps of ChIP-seq signal over 2kb regions centered on individual (C) hypomethylated nhDMRs and (D) hypermethylated nhDMRs ordered by percent methylation change. (E,F) Venn diagram of the number of intersections of H3K4me1, H3K4me3, and H3K27ac PTM marks with (E) hypermethylated nhDMRs and (F) hypomethylated nhDMRs. *** p < 0.001, ** p<0.01 using Fisher exact test.

### Hypomethylated nhDMRs are enriched for binding motifs of Nkx and other heart transcription factors

Several studies have shown DNA demethylation in intergenic regions of non-coding transcription can define active intergenic regulatory elements. Combining this with chromatin status can define functional classes of elements. Recent work comparing Nkx2.5 and p300 ChIP-seq data in mouse heart tissue identified Nkx2.5 as better predictor of cardiac enhancers (7). Nkx2.5 ChIP-seq studies have been carried out in immature and mature mouse cardiomyocyte cell lines, as well as mouse cadiac tissue (7,8,26,27). Several Nkx2.5 binding motifs have been proposed that show degeneracy in several base pairs amongst different studies. Zebrafish Nkx2.5 amino acid sequence closely resembles the human and mouse Nkx2.5 homologs, however, we find strong conservation of the homeodomain among all zebrafish Nkx family members, indicating all available Nkx motifs should be assessed (Figure S3A-B). To date no Nkx2.5 binding motif has been identified in zebrafish due to lack of availability of suitable ChIP-grade antibodies. To test for Nkx motifs in our subclasses of nhDMRs, we included all Nkx binding motif sequences from the Homer database, and loaded additional novel experimentally validated Nkx2.5 binding sites into the list of heart motifs. In addition, we compiled a list of all known Nkx2.5 binding partners from the literature (2,7,8,10,11,26) and extracted their known binding motifs from the Homer database (Figure S3C). We took all the filtered (>20% change in methylation) hypomethylated nhDMRs and used Homer to scan for enriched motifs. The predominant Nkx motif, observed in 1299 hypomethylated nhDMRs, had closer resemblance to Nkx6.1 motif than any other Nkx motif (Figure 4A). Since all Nkx family members share a conserved homeodomain, the most parsimonious explanation is that this is sequence of the more frequently used binding motifs for Nkx2.5 in zebrafish. Interestingly, for the hypermethylated class of nhDMRs, we observed a different Nkx motif with closer resemblance to Nkx2.1/2.5 motif and with a different combination of cardiac TF frequently co-clustering with this particular motif (Figure S4A-B).

**Figure 4:**
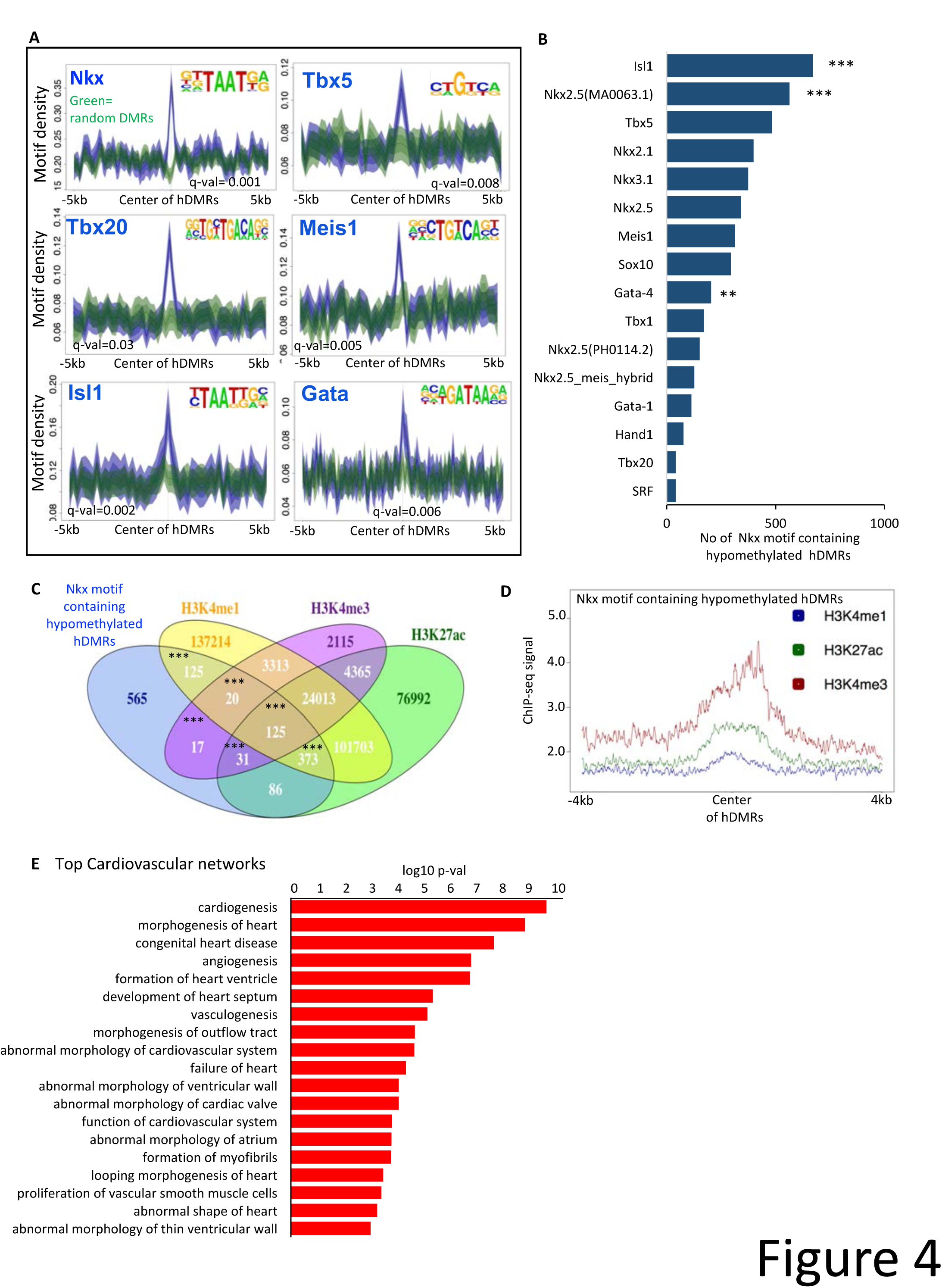
Hypomethylated nhDMRs associate strongly with Nkx family motifs, select cardiac TF motifs and with active histone PTM marks. (A) Homer analysis on hypomethylated nhDMRs identified enrichment of Nkx motifs and motifs for a subset of other heart transcription factors. Motif identity is indicated in top right hand corner and q-val in the bottom of panel. For a full list of Motifs enriched see Supplementary table S2. (B) Frequency of motifs for heart TF and Nkx family members found in significant Nkx-motif containing hypomethylated nhDMRs. Hypergeometric test, ** p-val < 0.01,***p-val < 0.001. (C) Venn diagram indicating number of hypomethylated nhDMRs that associate with histone PTM marks for active transcription enhancers. Fisher test, ** p-val < 0.01,***p-val < 0.001. (D) ChIP-seq signals of active histone PTM marks from 48hpf embryos were plotted over +/-4kb regions centered on hypomethylated nhDMRs. (E) GO categories of the signaling molecules enriched in the cardiovascular network. Expanded GO analysis of the cardiovascular network identifies genes integral to ventricle, atrium and valve formation.

Our Homer analysis found a striking number of significant cardiac TF motifs present among the Top 10 enriched motifs (Figure 4A, for full list see Table S2). We found Isl1, Gata-4 and other Nkx motifs to be significantly co-clustered with Nkx6.1 motif in the hypomethylated nhDMRs (Figure 4B). In contrast, Isl1,Tbx5 and Meis1 motifs co-clustered with the Nkx2.1/2.5 motif in hypermethylated nhDMRs. The presence of Isl1 in both categories of nhDMRS suggest a unique role for Isl1 in Nkx2.5-mediated heart development. Nkx2.5 and Isl1 positive cardiac progenitor cells give rise to multiple cardiac lineages. Although Nkx2.5 is shown to repress Isl1 in mESCs (28). Our data suggests a level of cooperativity between these transcription factors in regulating methylation of the nhDMRs. Further clustering of the nhDMRs by these distinct motif patterns shows evidence of Nkx2.5 dependendent co-regulation of different nhDMRs most likely in different subpopulations of cardiomyocytes (Figure S5A-B). Future work detailing the characterisation of the different clusters of nhDMRs *in-vivo* will yield greater insight into Nkx2.5 driven gene regulatory networks.

### Nkx motif-containing nhDMRs neighboring cardiac genes are associated with active histone marks

We next asked whether the Nkx motif nhDMRs can act as regulatory elements to the nearby genes. We intersected all 1299 Nkx motif-containing hypomethylated nhDMRs with annotated active histone ChIP-seq marks and found significant enrichment of H3K4me1, H3K4me3 and H3K27ac centered around the Nkx motif nhDMRs (Figure 4C,D). Interestingly, we also observed H3K4me1 and H3K27ac centered around Nkx motif hypermethylated nhDMRs although with markedly less association of H3K4me3 PTM (Figure S4C-D).

To evaluate the molecular pathways potentially regulated by the Nkx-motif hypomethylated nhDMRs, we examined the neighboring genes. Using Ingenuity Pathway Analysis (IPA)(29), we determined the pathways most enriched in the nearby genes. Two of the top networks were cardiovascular and muscle development (Figure S6A). Several unique pathways in cardiac development were enriched, including cardiac ventricle formation, looping morphogenesis and vascular smooth muscle formation (Figure 4E), strongly resonating with the *nkx2.5*^−/-^ zebrafish phenotype (3,4,30). Similar to the hypermethylated nhDMRs, we found cardiovascular system to be one of the major pathways associated with hypomethylated nhDMRs. Interestingly we observed a strong association with the endocardial genes, suggesting Nkx2.5 dependent gene regulation plays an important role in vascular signalling pathways (Figure S4E-F). We then tested if any of the Nkx motif hypomethylated nhDMRs were associated with altered expression levels of genes nearby. We found the Nkx intergenic nhDMRs had moderate but significant negative correlation with expression of the nearby genes, in particular, histone marks associated with intergenic and intronic regions (Figure S6B-C). This suggests Nkx motif hypomethylated nhDMRs exert regulatory roles on the expression of nearby genes.

### Nkx2.5 affects diverse signaling pathways during heart development

To determine whether nhDMRs contribute to the regulation of differential gene expression in Nkx2.5 mutants, we performed RNA-seq analysis on triplicate samples of *nkx2.5*^−/-^ and sib hearts at 48 hpf. We observed altered gene expression patterns of a moderate number of genes (Figure 5A,B). We found that a majority of the differentially expressed genes were upregulated (112 genes, adj p-value <0.05) versus downregulated (29 genes, adj p-value <0.05), suggesting *nkx2.5* is an activator of a small subset of genes but acts predominantly as a repressor. Genes that were downregulated included *vmhcl* (ventricular myosin heavy chain) and genes encoding calcium ion binding proteins and intermediate filament proteins, in agreement with the proposed function of Nkx2.5 in the maintenance of ventricle identity. Genes that were upregulated were enriched in several pathways including myofibril assembly, cardiac and smooth muscle contraction, consistent with the view that Nkx2.5+ progenitors cells can give rise to multiple cardiac cell lineages inlcuding conduction cell and smooth muscle cells (31). However these hypotheses are largely untested in zebrafish *nkx2.5*^−/-^ mutants. Interestingly, we observed a cohort of upregulated genes not characterized in the context of Nkx2.5 heart development, including mical2b, tjp-1b, c-fos, junba and c-jun. Mical2b is a nuclear actin regulatory protein, morphants of which show looping defects and reduced expression of cardiac and smooth actin proteins (32). Although Tjp-1b is expressed in the heart (33), its functional role has not been elucidated. C-fos and Junba, proto-oncogenes that are components of the AP-1 TF complex, are upregulated in response to cardiac hypertrophy and negatively regulate atrial natriuretic factor (ANF) (34), a target of Nkx2.5. C-jun has shown to be essential for the development of the outflow tract (35). Motif analysis on Nkx2.5 differentially expressed genes, did not identify any significant overrepresentation of Nkx motifs. This prompted us to investigate whether the differentially expressed genes showed strong correlation to the DNA methylation status of their TSS. Average fraction methylation profiles of differentially expressed genes were plotted and clustered by gene body DNA methylation from TSS to TSE (Figure S7A,B). We found no distinct differences between *nkx2.5*^−/-^ heart and sib heart in the overall gene body DNA methylation of these misexpressed genes. In addition gene expression was weakly correlated with respective DNA methylation level at their TSS (Figure S7C). This led us to ask whether neighboring nhDMRs could potentially regulate Nkx2.5-differentially expressed genes.

**Figure 5:**
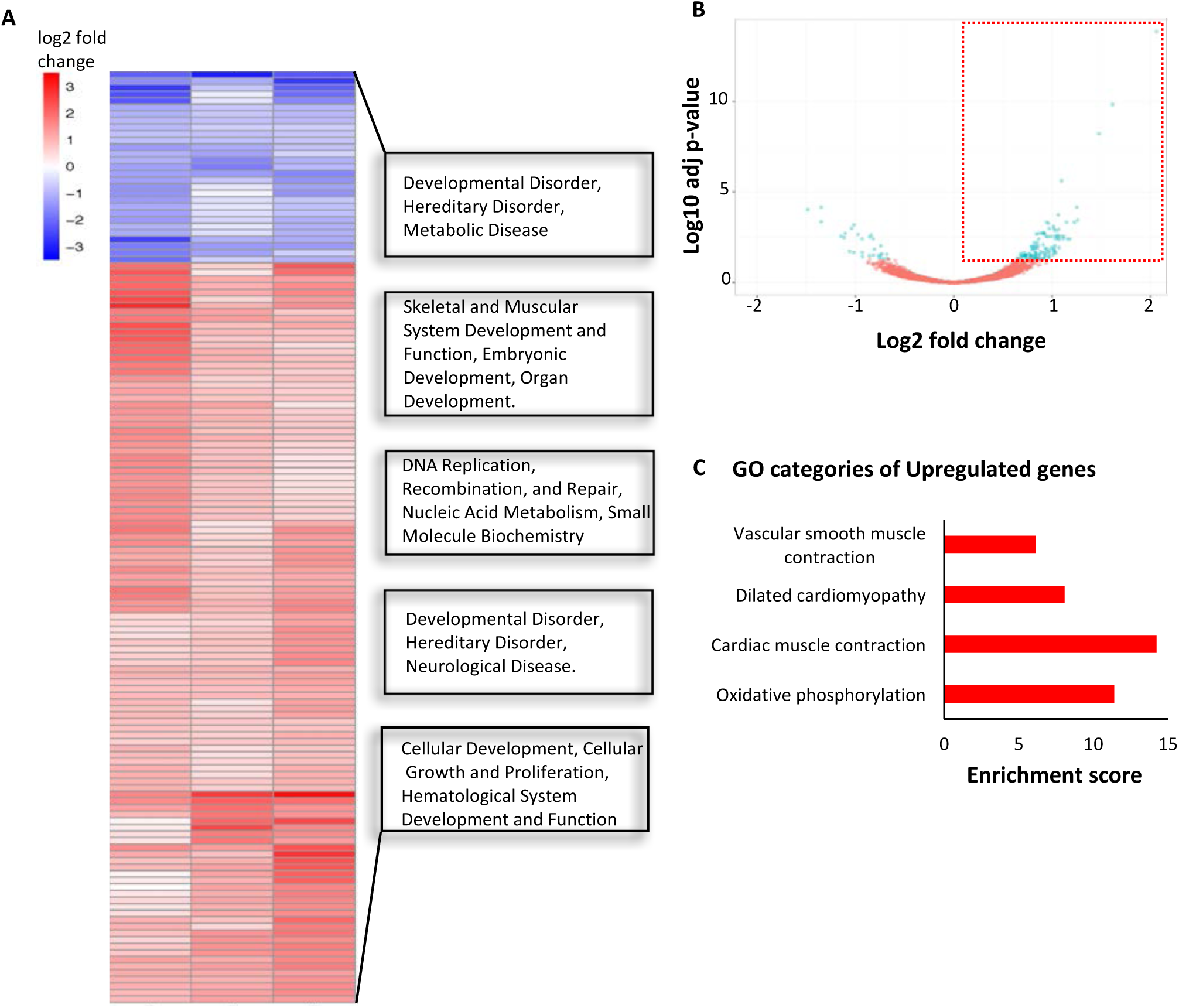
Vascular smooth and cardiac muscle contraction genes are strongly represented among differentially expressed genes in 48hpf *nkx2.5*^−/-^ heart. (A) Heat map of hierarchical clustering of differentially expressed genes. Genes in red denote upregulation and blue denotes downregulation in 48hpf *nkx2.5*^−/-^ hearts. GO clusters of differentially expressed genes identified by IPA analysis. Cardiac muscle contraction the most enriched GO pathways in gene upregulated in *nkx2.5*^−/-^ heart. (B) Volcano plot of *nkx2.5* differentially expressed genes. Red outlined box highlights the upregulated genes. (C) GO categories of genes upregulated in *nkx2.5*^−/-^ hearts, pertinent to nkx mutant phenotype.

### nhDMRs neighboring differentially expressed genes are enriched in Nkx motif

Nkx2.5 has been shown to bind and regulate genes via both proximal and distal promoters (36). In the developing mouse heart, selective expression of ANF is regulated by Nkx2.5 responsive cis-regulatory elements (36). To test if differentially expressed genes were near regulatory elements identified by our *nkx2.5*^−/-^ DNA methylome map, we took each nhDMR nearest to each differentially expressed gene and categorized it as a DMR in promoter, exon, intron or intergenic region. Of the 95 genes that are upregulated (excluding mitochondrial genes, since we observed no nhDMRs on mitochondrial genome), we identified 72 genes adjacent to a hypomethylated nhDMRs and 23 genes adjacent to an hypermethylated nhDMRs. Majority of the nhDMRs were intergenic. Using Homer, we found that Nkx(2.1/2.5) motifs were significantly enriched in a subset of the nhDMRs (Figure 6A, (p-val < 0.05)). Clustering of binding sites have successfully been used to scan the genome sequence to predict tissue specific enhancers; using these criteria we observed all Nkx motif-containing nhDMRs showed co-clustering of transcription factor binding motifs (Figure S8A). We calculated the average distance (bp) between Nkx enriched motif and other motifs in the nhDMRs. Using this list, we ranked the Nkx motif-containing nhDMRs in order of average distance (bp) from smallest to largest. We identified several other Nkx family members as the most frequently observed motifs that clustered with Nkx enriched motif (Figure S8B). This gave us a list of nhDMRs likely to be functionally responsive to Nkx2.5 regulation in-vivo.

**Figure 6:**
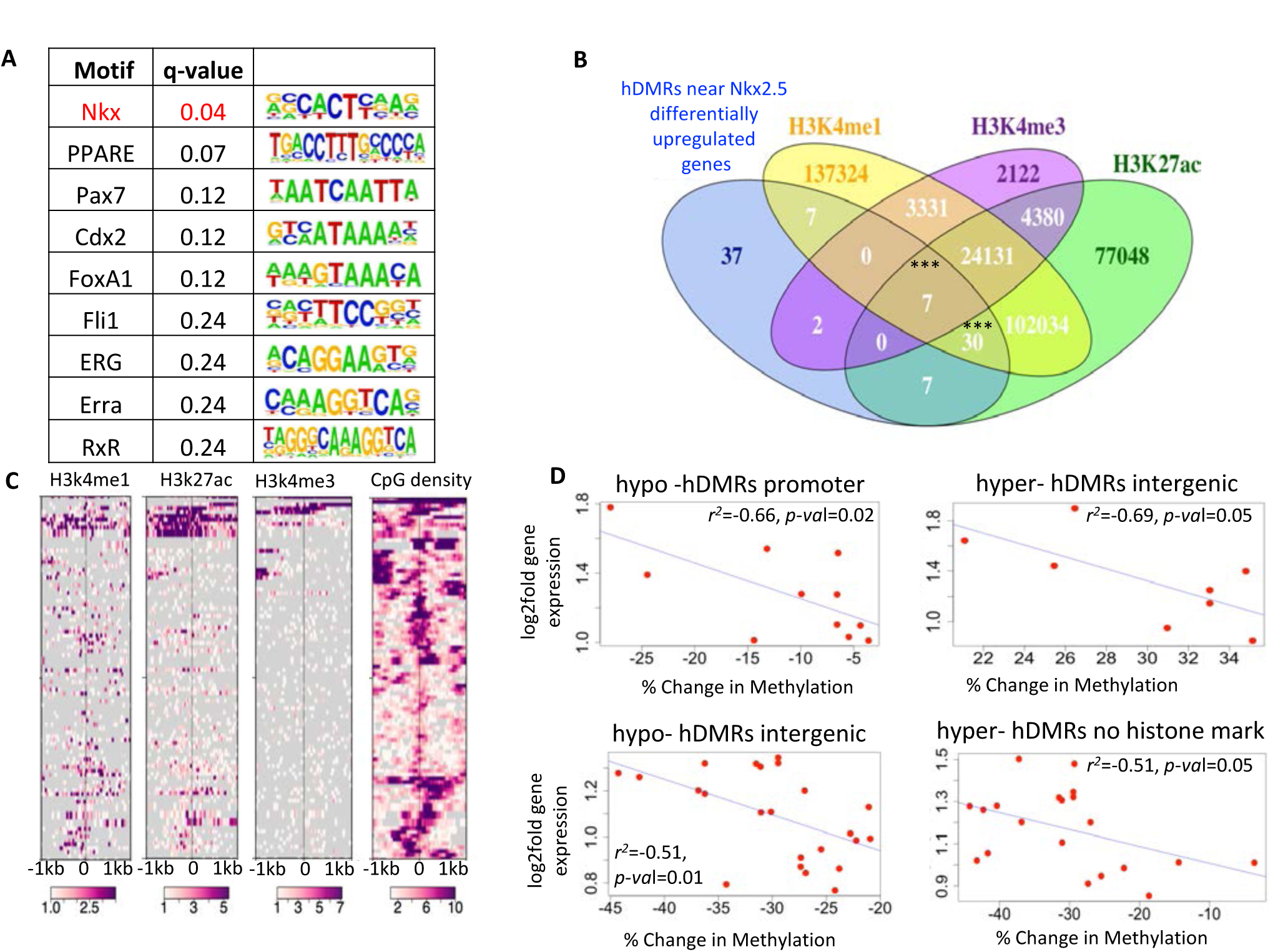
Nkx motif hypomethylated nhDMRs regulate nearby differentially expressed genes. (A) Top Motifs and consensus sequences observed in nhDMRs neighboring differentially expressed genes upregulated in *nkx2.5*^−/-^ hearts. Nkx motif was observed in 40 of the 95 nhDMRs. (B) Venn diagram intersecting the hDMRs nearest genes upregulated in *nkx2.5*^−/-^ hearts with the indicated H3K4me1, H3K4me3, and H3K27ac PTM marks. Fisher test, *** p-val < 0.001. (C) Heatmaps of ChIP-seq signal and CpG density mapped over 2kb regions centered on nhDMRs nearest to upregulated genes. (D) Percent change in methylation of nhDMRs plotted against the log2fold change in expression of the nearest differentially expressed gene.

To evaluate their functional potential, we identified a moderate number of nhDMRs that were significantly associated with either double (H3K4me1 and H3K27ac) or triple histone marks (H32k4me1, H3K4me3 and H3K27ac), with ChIP-seq signals enriched over the center of the nhDMRs (Figure 6B,C). To examine if the nhDMRs neighboring the upregulated genes showed potential regulatory function we looked for correlation with the differentially upregulated genes. Interestingly, we observed significant correlation with all promoter and intergenic nhDMRs with nearby gene expression (Figure 6D). Most notably, hypermethylated nhDMRs had strong correlation to nearby Nkx differentially expressed genes, suggesting a different mode of action for Nkx from that found in hypomethylated nhDMRs. Although we observed several strong negative correlations with the Nkx motif nhDMRs associated with a histone mark, these were not statistically significant due to the small number identified.

### Nkx motif nhDMRs regulatory elements validated in vivo

We sought to test whether the nhDMRs that we identified could function as enhancers of heart development. We selected nhDMRs near genes associated with heart development (*nkx2.5, cited2, ƒoxc1a*), genes encoding epigenenetic modification factors (*dnmt4,tet3*), and genes differentially expressed in the *nkx2.5*^−/-^ mutants from our transcriptome analysis (*ƒosb, myh7bb, tjp1b,smtnl1,myocd,cacna1*). We cloned these nhDMRs sequences into reporter constructs with a minimal promoter driving green fluorescent protein (EGFP) and injected zebrasfish embryos with the enhancer reporter vector along with Tol2 transposase messenger mRNA, to drive integration and stable transgenesis. Each group of G_0_ embryos injected with putative enhancer reporter constructs showed unique expression profiles in the embryo, with all showing expression in the heart. We established founder G_1_ transgenic fish that had offspring embryos expressing GFP, indicative of stable germline transgenics. We compared the GFP expression profiles to the endogenous whole mount in-situ expression of the genes documented in Zebrafish Model Organism Database (ZFIN), and found that the transgenic GFP expression patterns (Figure 7A-D) recapitulated the expression patterns of the endogenous candidate gene from which the neighboring nhDMR was derived. The seven nhDMRs we tested all matched their nearby candidate gene expression profiles. Although we paired each nhDMRs to its closest putative target gene, this does not rule out the possibility that the identified nhDMRs might regulate additional target genes whose patterns differ slightly from their endogenous gene expression. All nhDMR transgenics showed moderate to strong GFP expression in the heart, in particular the ventricle, suggesting that Nkx2.5 may play an essential role in regulating these elements in heart development. Although we tested a small subset of our identified nhDMRs, this approach paves the way for additional testing of putative nhDMR enhancer elements and their role in Nkx2.5 dependent heart development.

**Figure 7:**
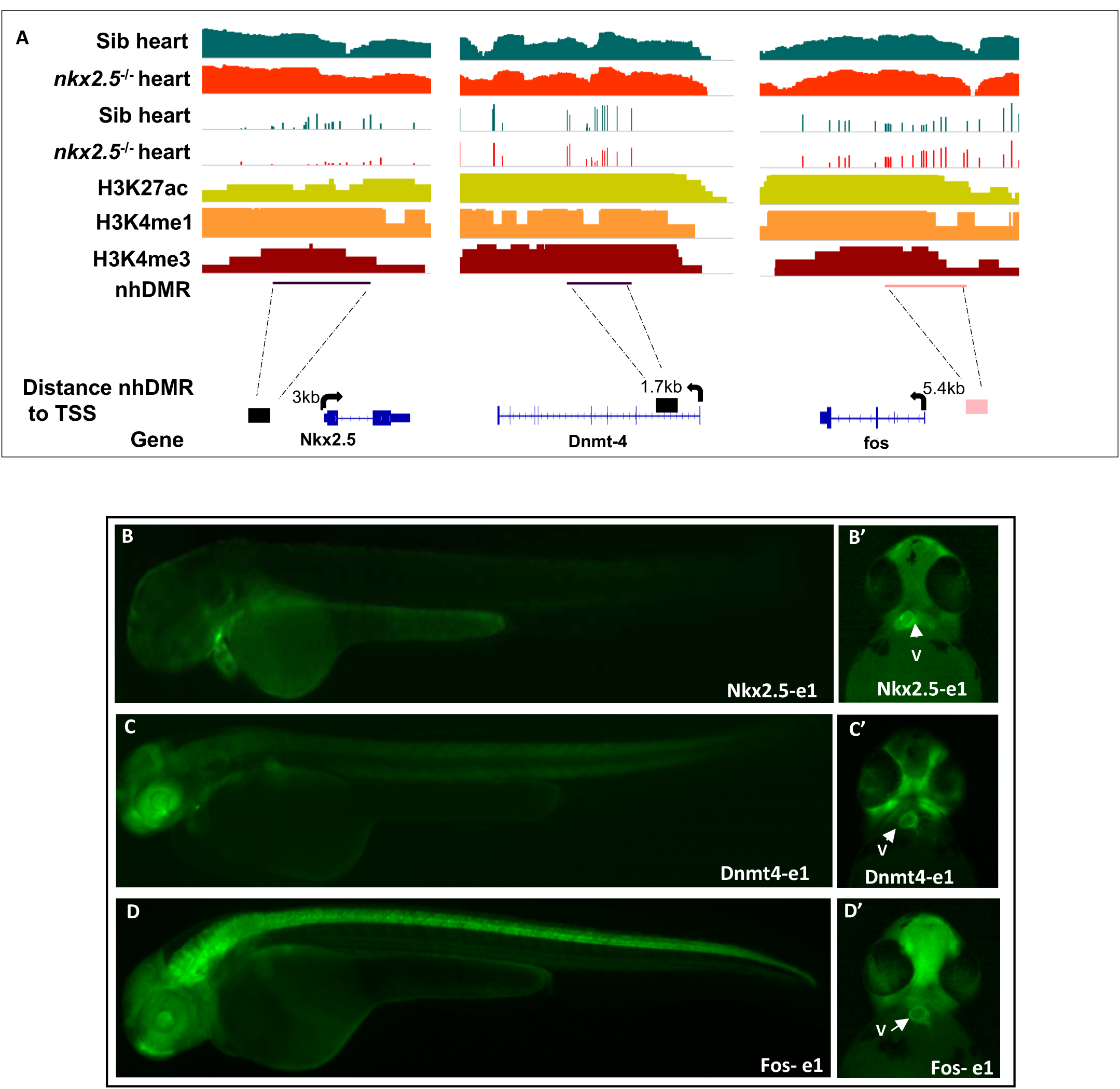
*In vivo* validation of Nkx putative enhancers. (A) Few examples of Nkx motif hypomethylated nhDMRs that serve as regulatory elements of nearby differentially expressed genes. First two rows from top denote DNA sequencing coverage in both samples. Third and fourth rows denote methylation tracks (5 reads or more per CpG) of sib and *nkx2.5*^−/-^ heart. Fifth through seventh rows indicate histone PTM ChiP-seq enrichment. Eight row indicates region of nearest Nkx motif hypomethylated nhDMRs. Ninth row lists calculated distance of nhDMRs to nearest differentially expressed gene and last row is a depiction of the direction of transcription of the gene. GFP expression driven by the the nhDMRs that we identified adjacent to the following genes: Nkx2.5, (B, B’) DNMT4, (C, C’) and Fos-b (D,D’). These nhDMRs appear to function as enhancer elements driving expression in the heart (B’-D’), stronger expression can be detected in the ventricle. V; ventricle, OFT; outflow tract.

## Discussion

DNA methylation is altered in *nkx2.5*^−/-^ mutant zebrafish embryonic hearts, indicating that Nkx2.5 is required for normal DNA methylation patterns in cardiac development. By generating comprehensive genome-wide DNA methylome maps at base-pair resolution in both wildtype and *nkx2.5*^−/-^ mutant zebrafish embryonic hearts, comparing these to DNA methylation patterns in non-heart embryonic tissues, we have identified novel Nkx2.5-dependent heart-specific Differentially Methylated Regions (nhDMRs). Integrating the dataset with the RNA-seq transcriptomes in mutant hearts, and wildtype histone PTM ChIP-seq data, we created a list of putative enhancers dependent on Nkx2.5 regulation in heart development. We further validated these *in vivo* by creating stable transgenic lines that show expression in heart.

Does Nkx2.5 have a direct role in regulating the methylation status of nhDMRs? The most parsimoniuous explanation is that Nkx2.5 directly binds nhDMRs containing Nkx motifs and participates either in the upregulatulation or downregulation of the methylation status of the nhDMRs to influence the expression of neighboring genes. In support of this explanation, it is striking that each class of nhDMRs, hypomethylated and hypermethylated, were enriched for distinct combinations of Nkx binding motifs and potential co-regulatory factors. Hypomethylated nhDMRs showed strong enrichment of the Nkx 6.1 motif and subset of other core heart transcription factors motifs (Gata4, Tbx5, Meis1, and Isl1). In contrast, hypermethylated nhDMRs were enriched for Nkx2.1/2.5 motifs. These observations suggest hypermethylated and hypomethylated nhDMRs have distinct modes of regulation by Nkx2.5 and cofactors. Importantly, both classes of nhDMRs were found to be conserved across vertebrate taxa, strongly suggesting widespread functional potential in cardiac development.

Our RNA-seq data corroborated earlier findings of Nkx2.5 roles in ventricle formation (3,4). We also observed that several genes important in heart development, including *bmp4, vcana, vmhcl and gata4*, were misregulated in *nkx2.5*^−/-^ hearts as previously reported. Intersection of the nhDMRs with ChIP-seq data identified significant association with H3K4me1 and H3K27ac PTM marks. Both marks are strongly linked to active regulatory elements, such as enhancers. Our transgenic assay provides strong evidence that nhDMRs function as enhancers in the heart and other tissues and show that changes in DNA methylation of the enhancer elements play a role in Nkx2.5 mediated heart development.

Since we observed a significant association of a subset of nhDMRs with unique Nkx binding motifs, we hypothesize that Nkx2.5 transcription factor has two roles: First, in the context of hypomethylated nhDMRs observed in *nkx2.5*^−/-^ mutants, we propose loss of Nkx2.5 acts to enhance expression of nearby differentially expressed genes most likely through a reduction in DNA methylation at the regulatory element (Figure 8A). Second, in the presence of Nkx2.5, the expression of the nearby gene is downregulated likely through increased DNA methylation of the regulatory element via the recruitment of cofactors (Figure 8B). Nkx2.5 comes from a family of homeobox proteins which can bind to methylated CpG via their homeodomain (19). So it is conceivable that Nkx2.5 could bind directly to CpG methylated DNA and foster the recruitment of co-factors to remove DNA methylation. In addition Nkx2.5 might also influence DNA methylation indirectly, by regulating the expression of DNA methylation machinery. Although Nkx2.5 has not been shown to directly bind at the promoter of HDAC1, we observe in multiple published Nkx2.5 ChIP-seq datasets (8,26) that Nkx2.5 binds at promoters of DNMT3a, TET1, TET2 and TET3 in several mouse derived cardiomyocyte cell lines. It is conceivable that Nkx2.5 binding events are amplified in a homogenous cardiomyocyte cell line which otherwise maybe masked in whole heart tissue experiment and makes it worthwhile investigating the DNA methylation profile of mouse derived cardiomyocyte cell line in response to Nkx2.5 knockout to further corroborate these hypotheses.

**Figure 8:**
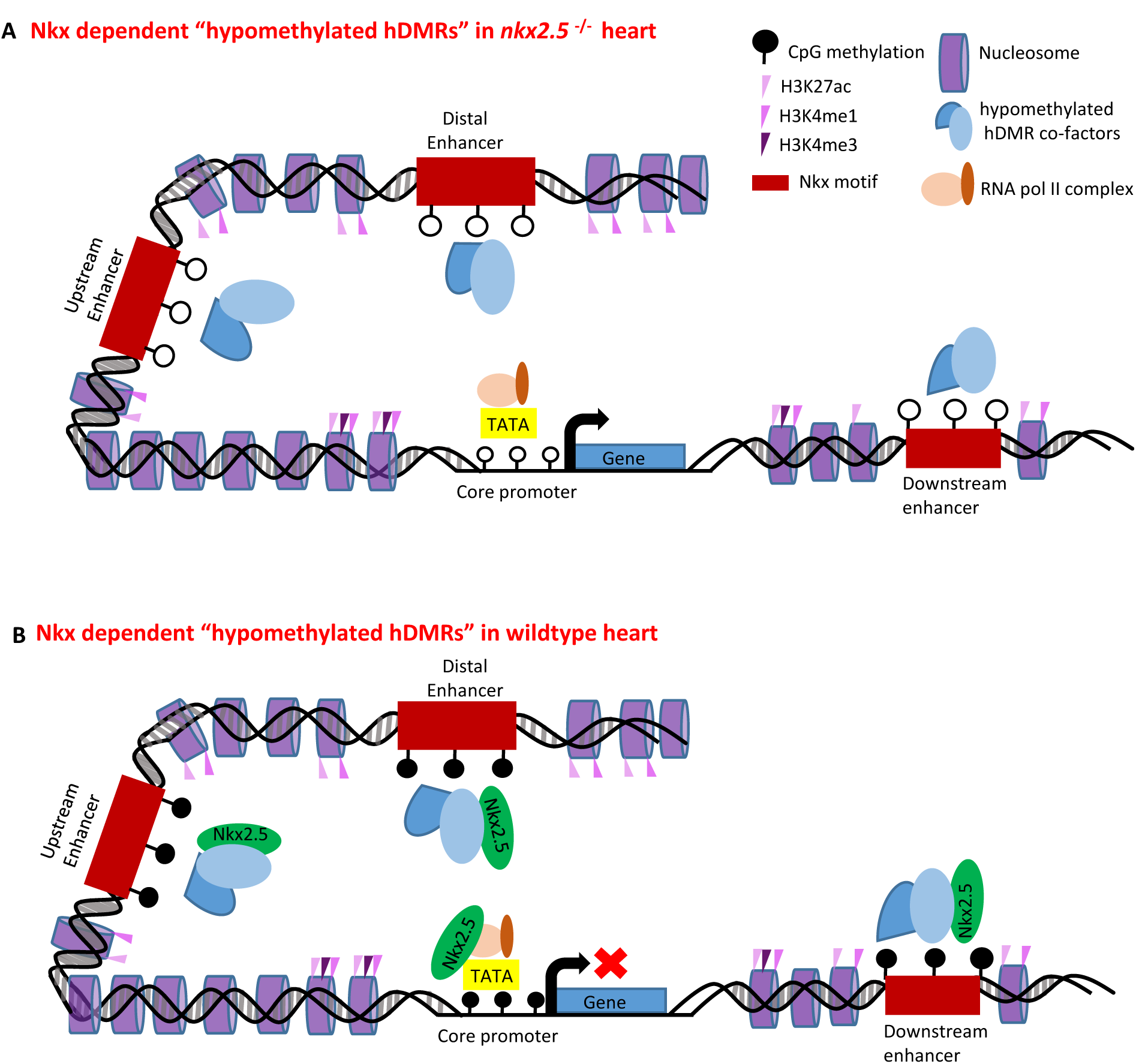
Proposed roles of Nkx2.5 in heart-specific DMRs neighboring differentially expressed genes. Hypomethylated nhDMRs containing Nkx motif are represented by red rectangular boxes located proximally (upstream and downstream) and distally relative to gene transcription start site. (A) In the context of hypomethylated nhDMRs observed in the *nkx2.5*^−/-^ mutant, we propose loss of Nkx2.5 acts to enhance expression of nearby differentially expressed gene. (B) In the presence of Nkx2.5 in wildtype embryos, the nhDMRs would increase in DNA methylation either through recruitment of “hypomethylated nhDMRs cofactors” by Nkx2.5 or by a direct role in allowing access of DNA methylation machinery.

When does Nkx2.5 influence the methylation status of nhDMRs? DNA methylation is highly dynamic during cardiomyocyte development, postnatal maturation and disease. In mammals, demethylated regions in neonatal and adult cardiomyocytes are localized in cell type-specific enhancer regions and gene bodies of cardiomyocyte genes (37). Previous characterization of *nkx2.5*^−/-^ zebrafish mutants suggested Nkx2.5 functions early, at 36hpf (5), Our RNA-seq datasets of sibling hearts detected high level of Nkx2.5 expression at 48hpf, consistent with our observation of strong enrichment of Nkx motifs in the nhDMRs. Furhermore, our recent Self Organizing Map (SOM) time course analysis of zebrafish wildtype heart development from linear heart tube (30 hpf) to fully looped heart (72 hpf) indicates that *nkx2.5* is expressed at steady state throughout these stages (Hill et al manuscript accepted). Since DNA methylation patterns can be inherited over several cell divisions, it is likely that the methylation status of nhDMRs is established by Nkx2.5 and cofactors early in development and that persistent methylation states in nhDMRs serve a critical function in maintenance of ventricle identity and downstream Nkx-mediated signaling pathways revealed by our analysis.

In summary, our results suggest that Nkx2.5 binding sites in the nhDMRs that we have identified participate in the methylation and demethylation of specific genomic regions. These observations provide a platform for further investigation of heart transcription factors and their participation in the regulation of DNA demethylation. The different transcription factor binding motifs identified in conjunction with Nkx motifs within subsets of hypermethylated and hypomethylated nhDMRs suggests interaction with different cardiac TFs in the regulation of DNA methylation of nhDMRs. This model provides an explanation for the dual nature of Nkx2.5 to act as an activator and repressor during cardiac development. Our study sets the foundation for addressing the functions of Nkx-dependent elements in the heart and to uncover Nkx driven gene regulatory networks in the heart. These nhDMRs co-localize with gene regulatory elements, particularly enhancers and transcription factor-binding sites, which will allow identification of key lineage-specific regulators. Therefore, the highly conserved Nkx motif nhDMRs that we have identified can serve as a starting point to explore new putative regulatory elements in congenital heart disease.

## Methods and Materials

### Zebrafish stocks

Adult *nkx2.5*^+/-^ were outcrossed to *cmlc2*: GFP fish maintained on AB background. Adult carriers for nkx2.5 mutations were genotyped by High Resolution Melt Analysis (HRMA) (38) with the following primers: Fwd Primer; CAAACTCACCTCCACACAGG, Rev Primer; GCTGCCTCTTGCACTTGTATC. All adult fish were housed in the University of Utah Centralized Zebrafish Animal Resource facility. Animal research protocol 15-06004 was approved by University of Utah IACUC. Embryos were raised under standard conditions at 28°C. Embryos raised from (*nkx2.5*^+/-^ *; cmlc2*: eGFP) crosses were screened at 48hpf when the *nkx2.5*^−/-^ mutant phenotype is identifiable with an enlarged atrium and small ventricle. *nkx2.5*^−/-^ mutants were segregated from siblings that consisted of mixed genotypes (*nkx2.5*^+/+^ and *nkx2.5*^+/^). This nkx2.5 allele has a stop codon in the homeodomain, and heterozytotes do not have a discernable cardiac phenotype. Genotypes were subsequently confirmed in RNA-Seq and whole genome Bisulfite-sequencing datasets.

### *nkx2.5*^−/-^ heart collections and RNA-sequencing

Zebrafish embryos were first segregated by phenotype into *nkx2.5*^−/-^ mutants and sibs at 48 hours post fertilization (hpf) and then hearts were isolated; 150 hearts were collected per biological replicate for each genotype, using a protocol modified from previously published zebrafish heart isolation techniques (39). Isolated phenotyped hearts were washed twice with heart media and hand selected to further separate hearts from non-heart tissue and debris. Heart tissue was snap frozen in liquid nitrogen until sufficient samples were collected. RNA was purified from heart tissue following the QIAGEN Microplus Easy kit protocol. RNA samples were pooled together per replicate in order to collect an optimal amount for library preparation and sequencing. RNA quality and quantities were confirmed on the Bioanalyser RNA 6000 Pico Chip. cDNA libraries were prepared using the Illumina TruSeq kit at the University of Utah High Throughput Genomics Core Facility. All six samples, consisting of three matching *nkx2.5*^−/-^ mutants and sibling pairs were run on a single lane and 50bp single end reads were generated on the HiSeq 2000 machine. Samples were aligned to zebrafish Zv9 Ensembl version 77 genome annotation build using Novoalign with default parameters.

### RNA-seq analysis

We used DESeq2 default settings to measure differential expression between *nkx2.5*^−/^ heart and sib hearts, controlling for collection time. Gene read counts were generated using USeq “Defined Region Differential Seq” from the USeq package (40).

### Bisulfite-sequencing and analysis

Hearts and non-heart tissue were isolated as described above *nkx2.5*^−/-^ mutants and siblings at 48hpf, and genomic DNA was extracted from each genotype. A total of 100ng was collected from 400 hearts, and corresponding non-heart tissue, using the Qiagen non-organic DNA clean up kit. DNA samples were collected in triplicate, sheared using COVARIS, and spiked with 1% unmethylated lambda. Samples underwent bisulfite treatment using Epitect bisulfite kit and a total of 12 samples were each sequenced on four lanes of HiSeq 2000 to generate 101bp paired end reads. Fastq files for each replicate per genotype and tissue were merged from each run to give one final fastq file per replicate, and aligned to Zv9 using Novoalign. Each replicate per condition had 3x-5x median coverage, and genotypes were confirmed for each sample. Bioinformatics analysis on the Genome wide Bisulfite sequencing used tools from the USeq package (http://useq.sourceforge.net) (40) to calculate differentially methylated regions on merged replicates from each genotype. Each merged replicate had 9x-15x coverage minimum. To test if merged samples provided good representation of the three replicates, we ran PCA on differentially hypomethylated and hypermethylated regions in each condition. Fraction methylation of defined regions in individual replicates clustered close to the merged replicate (averaged fraction methylation)(Figure S1A-B).

## Analysis of nhDMRs

### Motif analysis and binding site prediction

Location of transcription factor binding sites and motif enrichments within DMRs were determined using default parameters in Homer (41). Additional Nkx2.5 motifs from recent published datasets (7,8,26) were also incorporated into the search. 50,000 random background sequences normalized for CpG content were chosen for the different subclasses of nhDMRs tested via Homer settings.

### Cis-regulatory element analysis (CEAS), evolutionary conservation and GO analyses

The CEAS script build_genomeBG was used to create a custom CEAS annotation file(42). Coverage data was used from all four conditions: *nkx2.5*^−/-^ heart, sib heart, *nkx2.5*^−/-^ body and sib body. Bed files of Differentially Methylated Regions (DMRs) were loaded into the program. Genomic coordinates were downloaded from UCSC zv9 ref gene annotations. For evolutionary conservation of nhDMRs the fish and vertebrate PhastCons score based on the eight by eight way fish and vertebrate genome alignment was obtained from the UCSC Genomer Browser PhastCons conservation track (https://genome.ucsc.edu/). The data was generated by averaging phastcon scores across the DMRs and counting the number of DMRs that have average phastcon score of > 0.5. randomBed was used to create 10,000 random regions of matched size of each DMR. mapBed was used to calculate average phastcon scores across these random regions. P-value was calculated by testing how many of the random regions have more conserved regions than defined DMRs. Gene ontology (GO) analysis of the neighboring genes of nhDMRs was carried out using Igenuity pathway analysis (IPA)(29) and DAVID (https://david.ncifcrf.gov/).

### ChIP-seq analysis

Histone ChIP-seq data of 48hpf whole zebrafish embryos was retrieved from Bogdanovic *et al* 2009 (22) and re-processed using MACs 2.0 software with default settings (43). DMRs that intersected with all three histone peak H3Kme1, H3K4me3 and H3K27ac independently of each other were merged together as one single peak for identification.

### Testing cis-regulatory elements in vivo

Eight nhDMR sequences with putative regulatory activity (S4 table) were PCR amplified with specific primers and TOPO-cloned into entry vector pCR8/GW/TOPO of the Gateway cloning system (Invitrogen). The cloned plasmid was then recombined with the destination vector pGW_cfosEGFP (21) to generate the desired reporter plasmid. Tol2 transposase mRNA was transcribed in vitro from pcs2+ using the mMessage mMachine Sp6 kit (Ambion). Embryos at one-cell or two-cell stages were injected with transposase mRNA. The reporter expression patterns were analysed at 24 hpf stage. If 10% of embryos exhibit the consistent GFP expression pattern, the construct was considered as a positive enhancer. The embryos with specific expression patterns were selected and raised to sexual maturity. Sexually mature G0 adults were crossed with AB wild-type strain to obtain germline transmission. Three independent F1 transgenic lines were established for each construct, unless otherwise indicated. F1 embryos were photographed at 48hpf stage.

## Declarations

### Data Access

All the data described in this paper can be downloaded from https://b2b.hci.utah.edu/gnomex/ RNA-seq experiment ID 246R, and Bisulfite seq experiment ID 324R.

### Ethics Statement

All genome sequencing data are fully available without restriction, as indicated above.

### Animal Research Protocols

Zebrafish animal research were reviewed and approved by the University of Utah IACUC, protocol number 15-06004.

### Funding and Acknowledgements

BG was supported by AHA Postdoctoral Fellowship (12POST12030301). This study was funded by a NHLBI Bench-to-Bassinet Consortium (http://www.benchtobassinet.com) grant to HJY (2UM1HL098160) and sequencing was provided by a core facilities support grant to New England Research Institute (U01 HL098188). We thank Bradley Demarest and Darren Ames for performing Bisulfite seq alignments, Kimara Targoff for discussions and supplying *nkx2.5*^−/-^ mutants and David Nix for bioinformatics discussions. The funders had no role in study design, data collection and analysis, decision to publish, or preparation of the manuscript.

## Supporting Information

**Figure Supplement 1:**
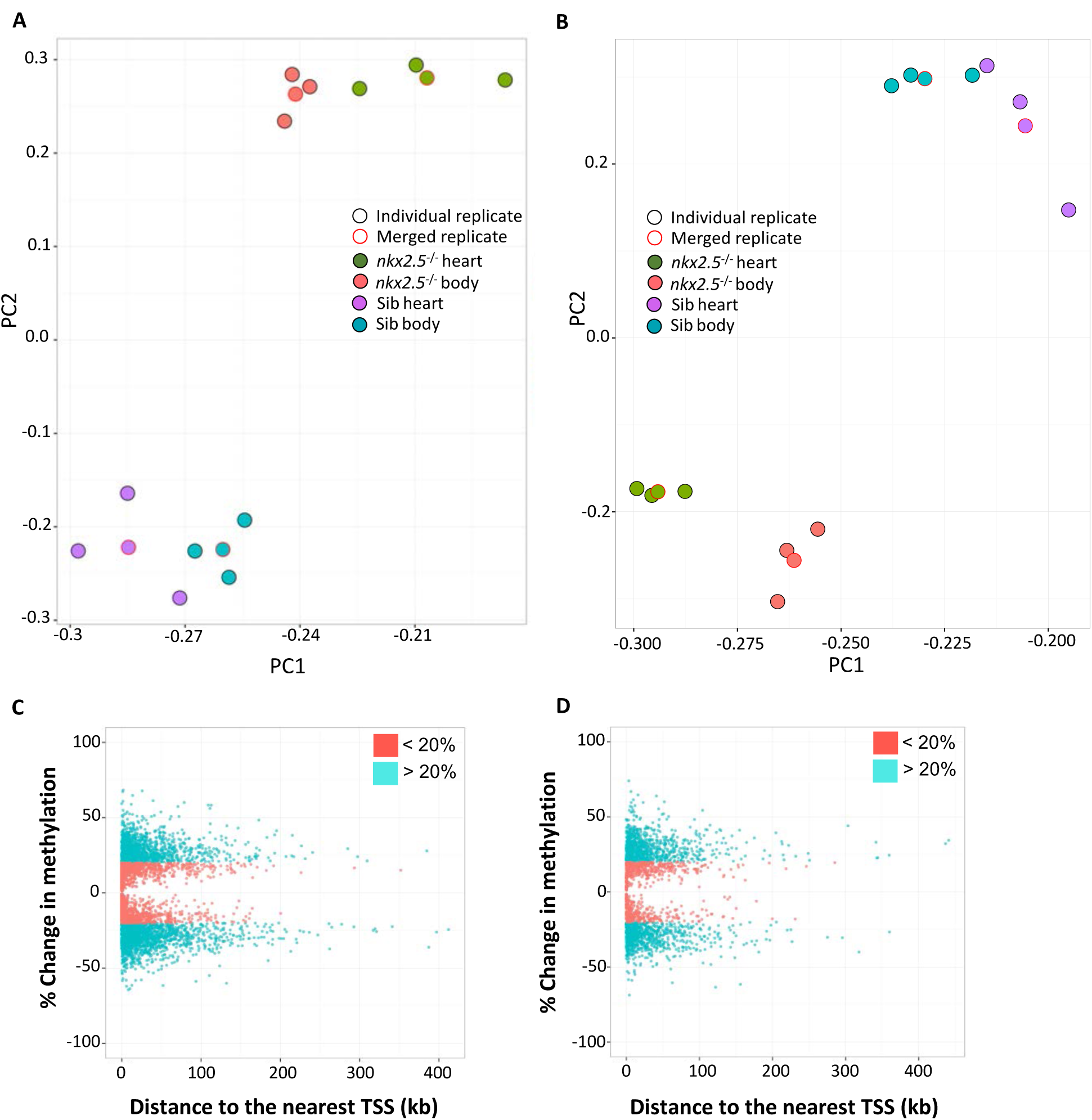
Clustering of individual replicates per condition and percent methylation changes in Nkx and Sib nhDMRs compared to non-heart tissue. PCA plots on the Fraction methylation of (A) hypomethylated DMR and (B) hypermethylated DMRs in each replicate (black outline circle) and averaged fraction methylation in merged sample (red outline circle). The first two PCA components were plotted. (C) Percent changes in methylation of DMRs from *nkx2.5*^−/-^ heart v. *nkx2.5*^−/-^ body plotted against distance to the nearest TSS. (D) Percent changes in methylation of DMRs from sib heart v. sib body plotted against distance to the nearest TSS. Less than 20% change is indicated in orange/pink and greater than 20% in cyan.

**Figure Supplement 2:**
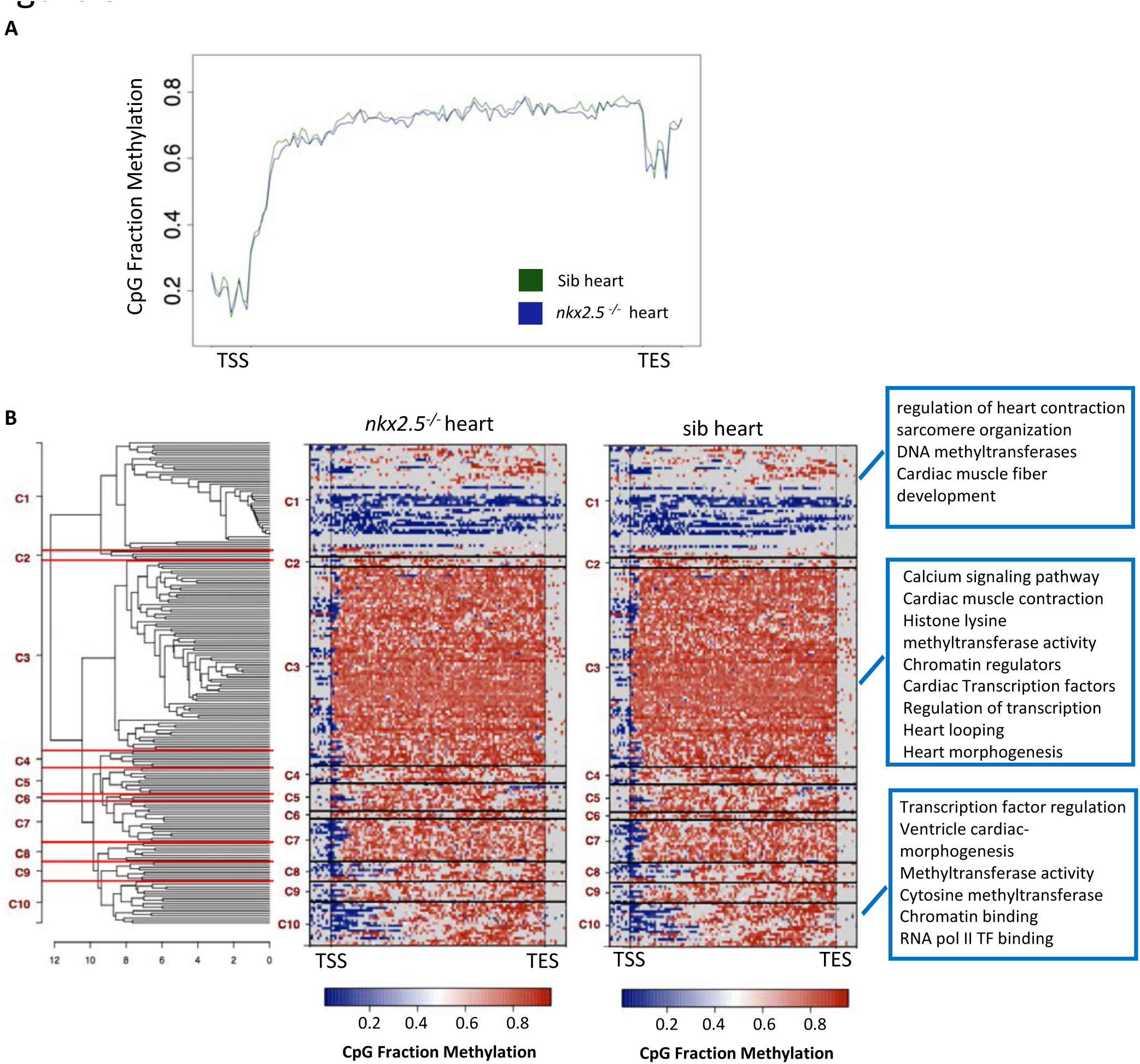
Gene body methylation of core cardiac and epigenetic markers in *nkx2.5*^−/-^ heart and sib heart. (A) Average fraction methylation summed over entire gene body lengths (normalized) and 100bp beyond each end of gene body. A complete list of core cardiac and epigenetic markers is in the S3 file. (B) Hierarchical clustering of core cardiac and epigenetic markers based on average fraction methylation from TSS to TSE. Blue represents regions of low CpG methylation and orange high CpG methylation. GO categories of genes in several clusters are listed in blue boxes.

**Figure Supplement 3:**
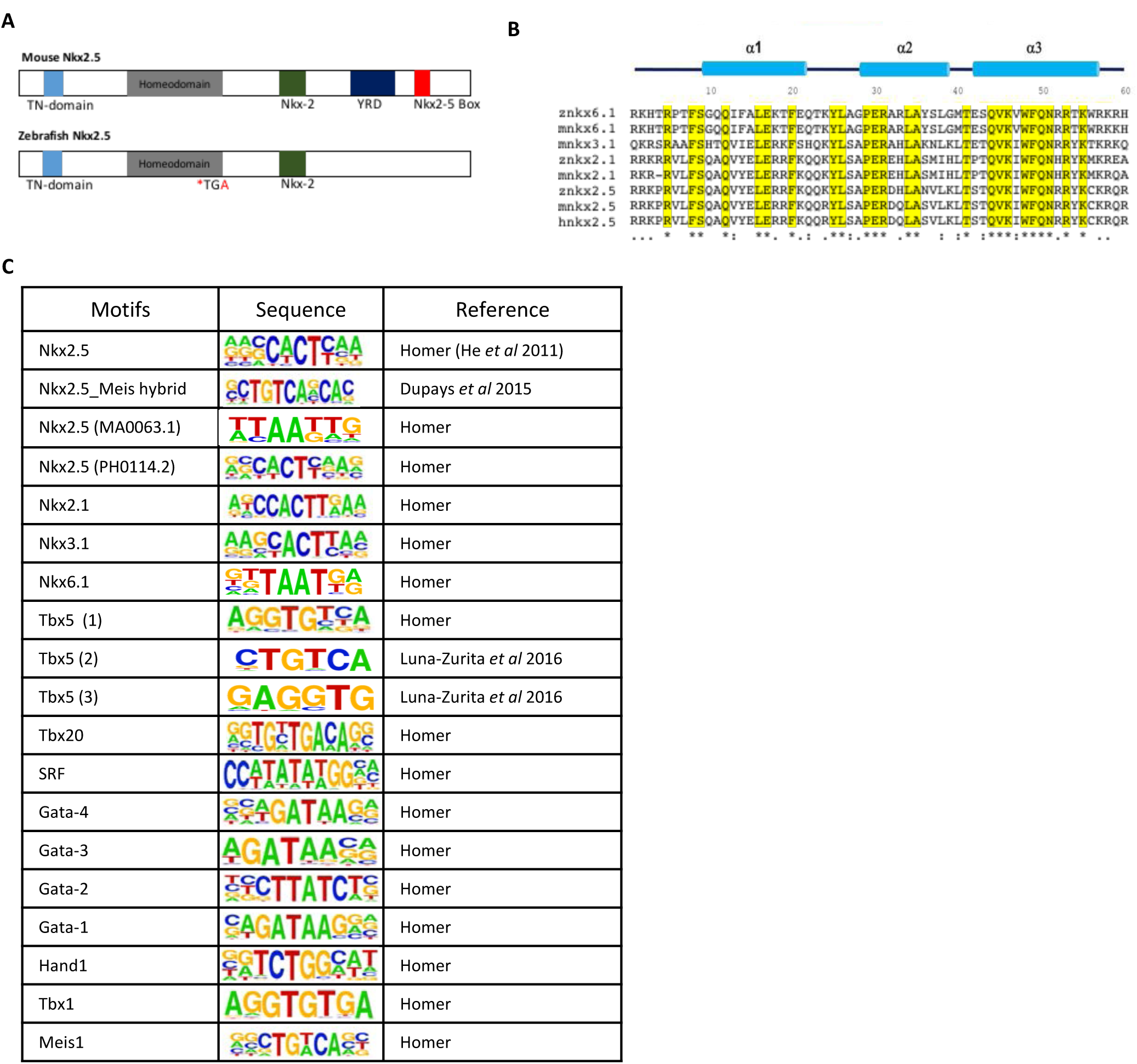
Nkx2.5 protein structural domains and homeodomain. (A) A schematic of zebrafish structural protein domains compared to mouse Nkx2.5 protein. YRD domain and carboxy-terminal Nkx2.5 box are missing in zebrafish. *TGA indicates site of mutation in zebrafish homeodomain. (B) Amino acid sequence of zebrafish, mouse and human Nkx family members homeodomains. Yellow indicates conserved amino acids. (C) Transcription factor binding motifs used in Homer analysis for TFs known to interact with Nkx2.5 and all zebrafish Nkx family members. References indicate sources for motif sequences.

**Figure Supplement 4:**
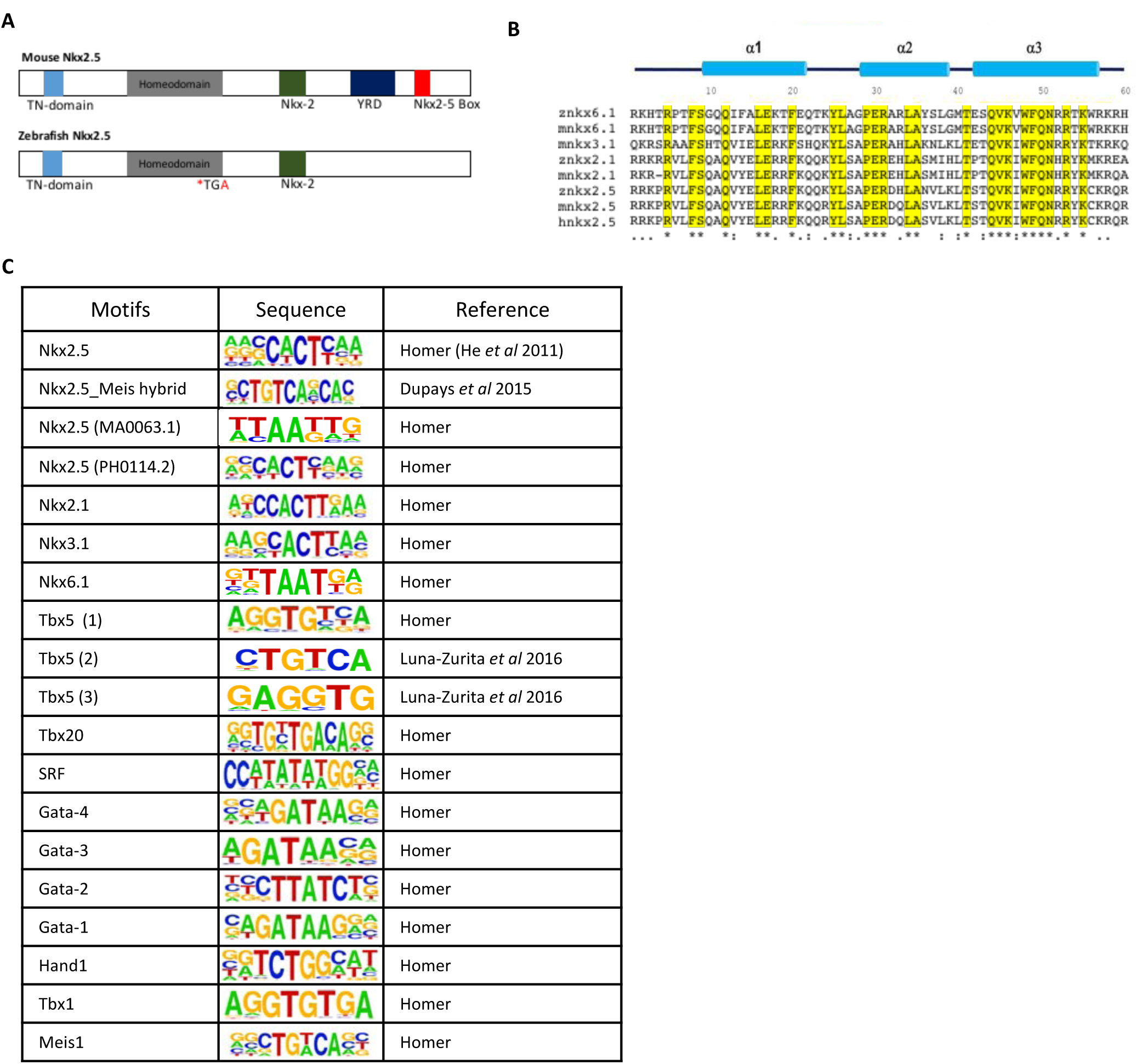
Hypermethylated nhDMRs associate strongly with Nkx family member motifs and with active histone PTM marks. (A) Homer analysis on hypermethylated nhDMRs identified enrichment of Nkx motifs and other heart TF motifs. Significant enriched motif is indicated in the top right hand corner and q-val in the bottom of each panel. For a full list of Motifs enriched see Supplementary table S2. (B) Frequency of heart TF motifs and other Nkx family member motifs found in significant Nkx motif hypermethylated nhDMRs. Hypergeometric test,* p-val<0.05, ** p-val < 0.01,***p-val < 0.001. (C) Venn diagram of number of Nkx motif hypermethylated nhDMRs associated with histone PTM marks associated with active enhancers, Fisher test, ** p-val < 0.01,***p-val < 0.001. (D) ChIP-seq signals of active histone PTM marks from 48hpf embryos were plotted over +/-4kb regions centered on Nkx motif hypermethylated nhDMRs. IPA analysis of the neighboring genes of Nkx motif hypermethylated nhDMRs. (E) Top networks associated with neighboring genes in hypermethylated DMRs. (F) Expansion of one of the top networks associated with cardiovascular function.

**Figure Supplement 5:**
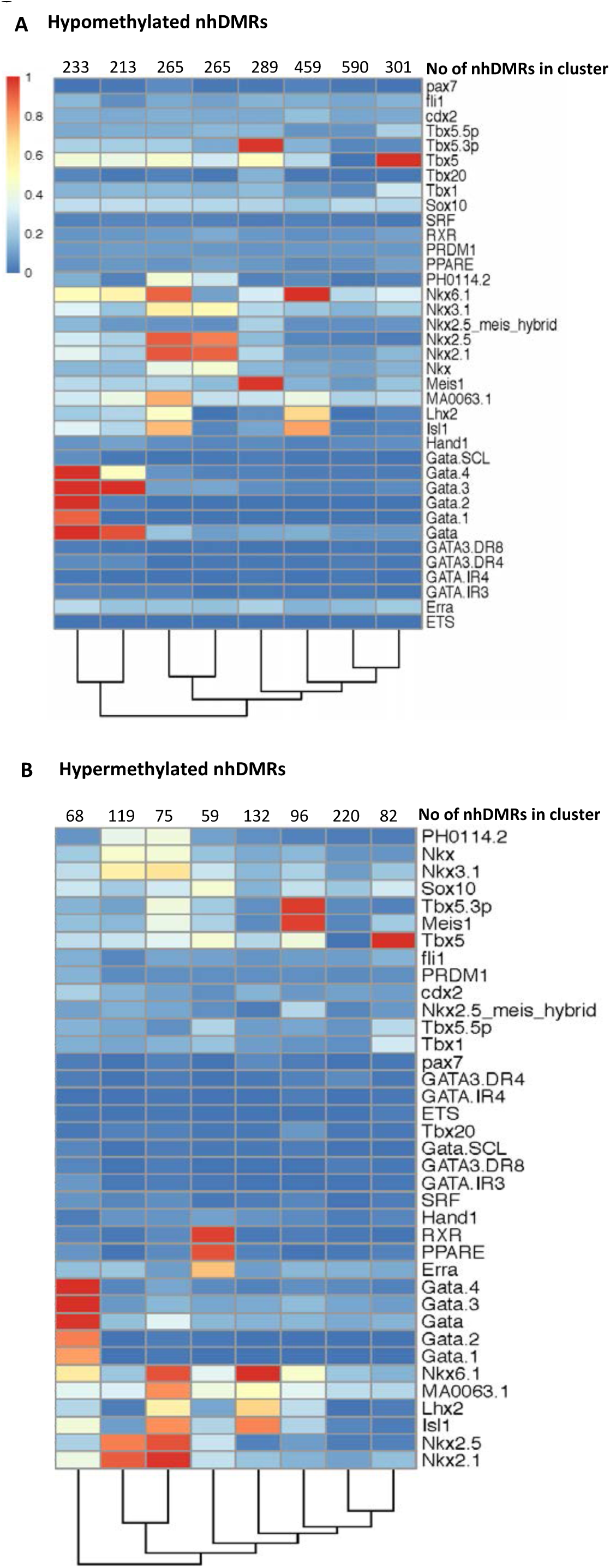
Distinct clusters of hypomethylated and hypermethylated nhDMRs based on motif patterns. K-means clustering on the motif patterns of hypomethylated nhDMRs (A) and hypermethylated nhDMRs (B). Each column is a cluster of nhDMRs, where the values is the average occurrence for a motif. A value of 1 (red) means all the nhDMRs in that cluster have that motif, while 0 (blue) means none of the nhDMRs have that motif. The motifs are in represented in rows. The number of clusters (k) being 8.

**Figure Supplement 6:**
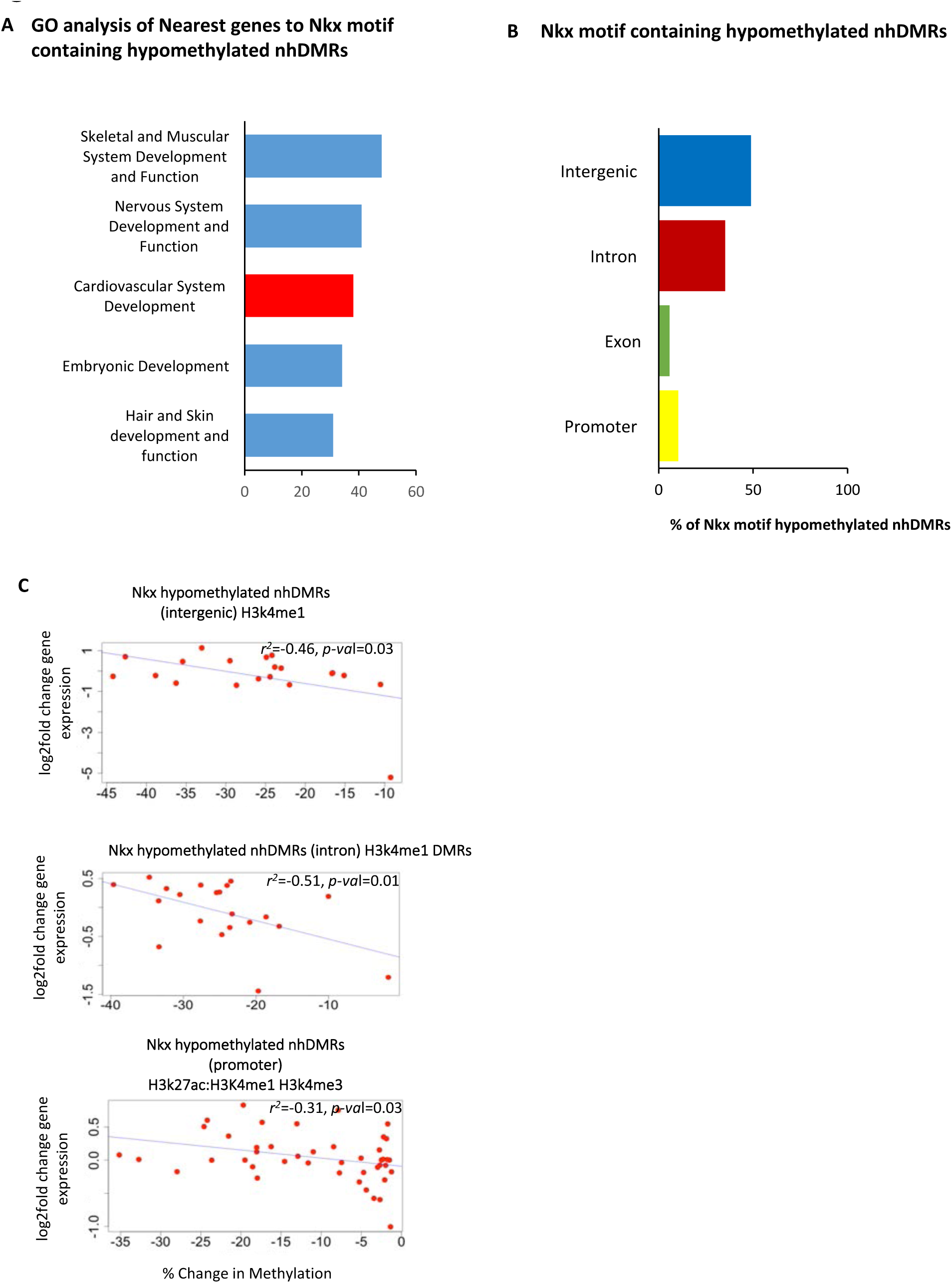
Nkx motif containing hypomethylated nhDMRs negatively regulate expression of neighboring cardiac genes. (A) Top networks represented with cardiovascular development being among the top networks in this subset of genes. (B) Categorization of the Nkx motif containing hypomethylated nhDMRs categorized into different genomic regions. Nkx motif hypomethylated nhDMRs are predominantly located in intergenic regions. (C) Genomic regions of Nkx motif hypomethylated nhDMRs and their correlation with nearest gene expression. Pearson correlation and p-value is reported on plot.

**Figure Supplement 7:**
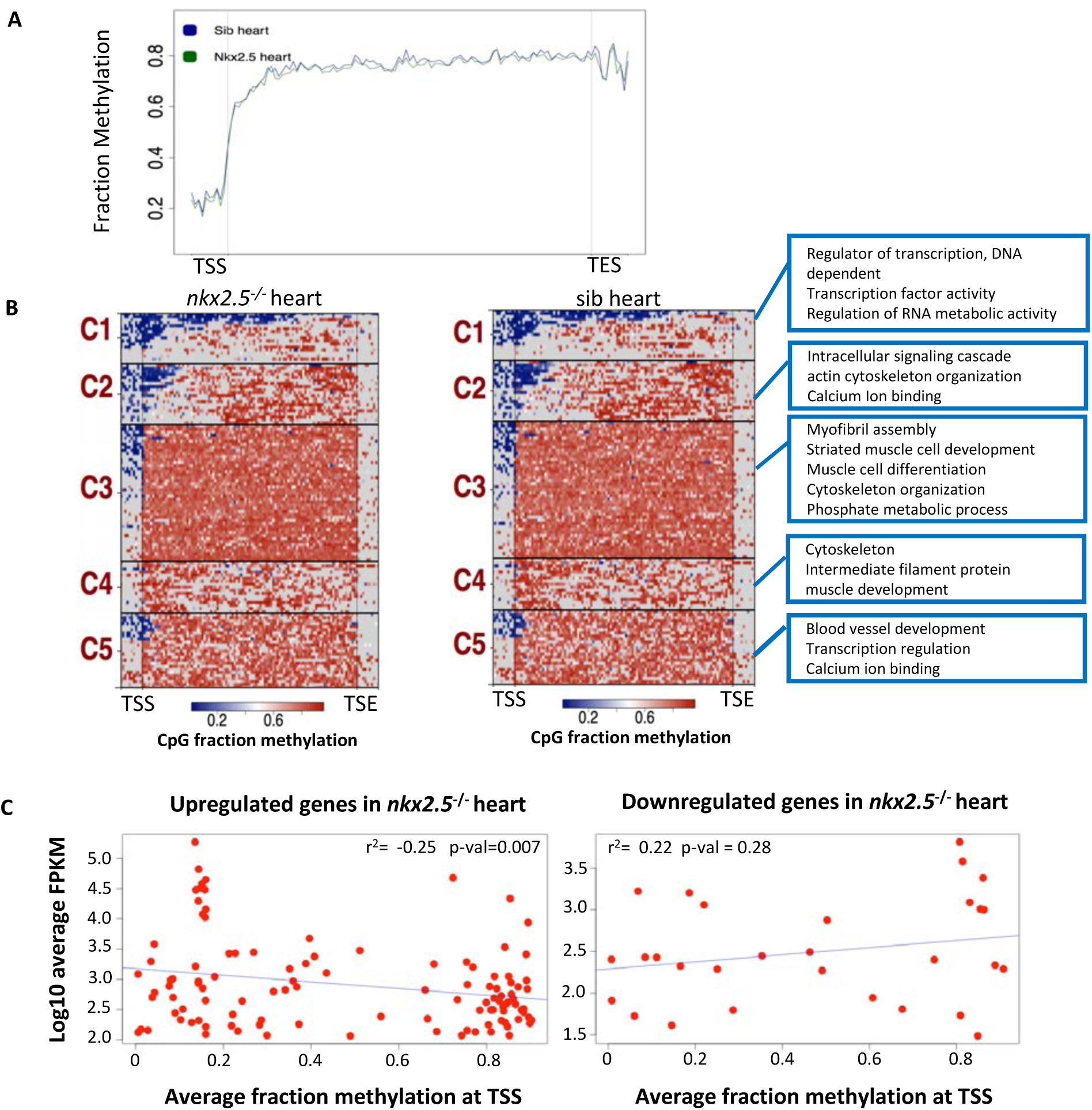
Gene body methylation of the differentially expressed genes in *nkx2.5*^−/-^ hearts. (A) Average fraction methylation summed over entire gene body lengths (normalized) and 100bp beyond each end of gene body. TSS and TSE denote transcription start and end. (B) K-means clustering of differentially expressed genes based on average fraction methylation across gene body. Blue represents regions of low CpG methylation and orange high CpG methylation. GO categories listed on the right of heatmaps. (C) Negative correlation of TSS methylation of differentially upregulated genes and their expression in *nkx2.5*^−/-^ heart. Positive correlation of TSS methylation of differentially downregulated genes and their expression in *nkx2.5*^−/-^ heart.

**Figure Supplement 8:**
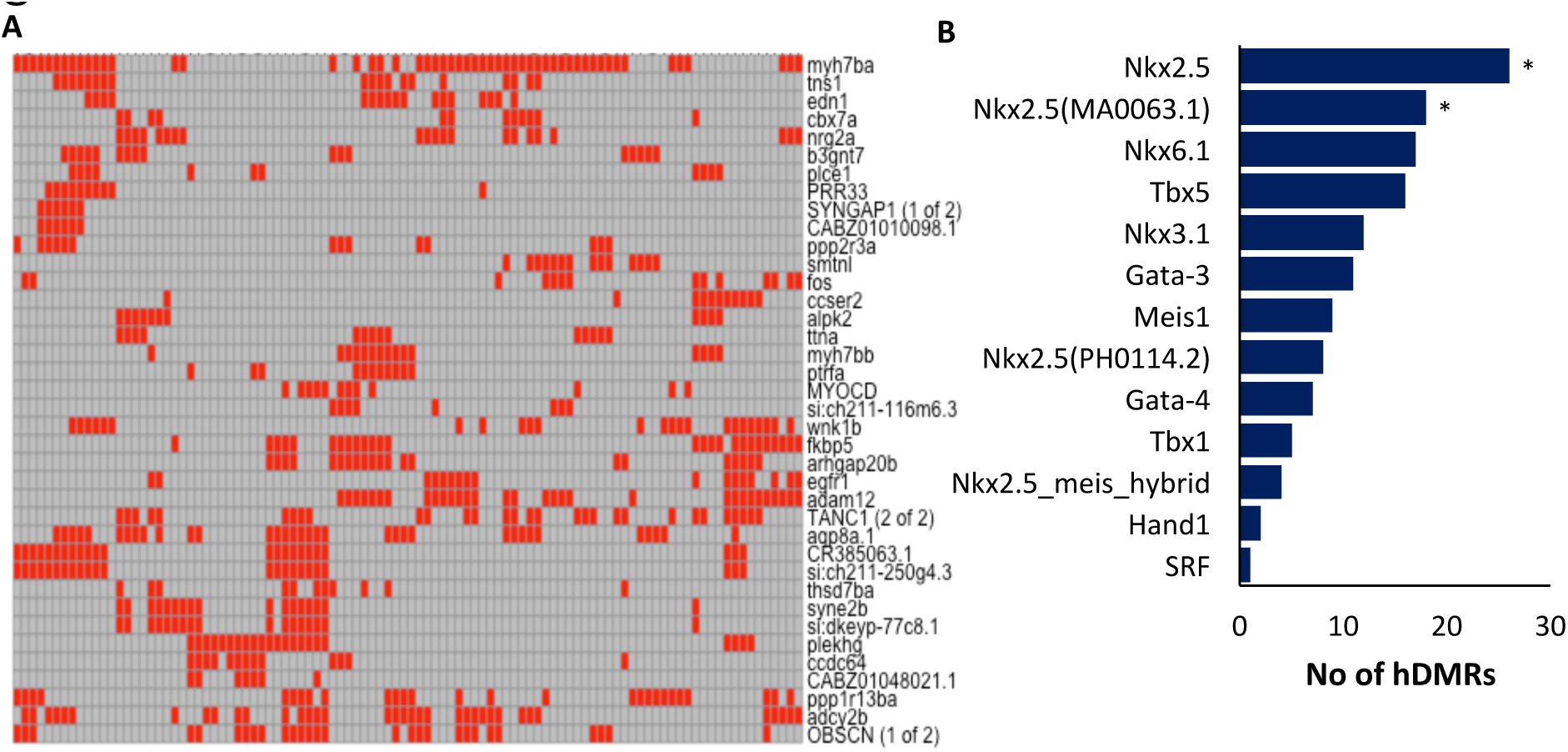
Nkx motif patterns in nhDMRs neighboring genes upregulated in *nkx2.5* mutants. (A) Heatmap of the motifs arrangement in each of nhDMRs neighboring each differentially upregulated gene in nkx2.5 mutant hearts. All nhDMRs were scaled to 100bp and average distance between motifs was calculated per nhDMRs and scored in a table and plotted on a heatmap (B) Most frequently associated motifs in the Nkx containing nhDMRs *p < 0.05.

